# African origin and Late Cretaceous divergence of the Middle American catfish *Lacantunia enigmatica* corroborated by a global mitogenome phylogeny

**DOI:** 10.64898/2026.01.30.702944

**Authors:** Jairo Arroyave, Adán Fernando Mar-Silva, Alfonso A. González-Díaz, Sebastián Huacuja-Barraza, Christian Lambarri-Martínez, Nelson R. Salinas, Kritsada Katawutpoonphan, Rupert A. Collins, Julia J. Day

## Abstract

Two decades ago, the new siluriform family Lacantuniidae was erected to accommodate its sole known extant representative, *Lacantunia enigmatica*, a morphologically aberrant catfish restricted to the Middle Usumacinta River basin along the Guatemala-Mexico border. While its discovery was unexpected, its proposed phylogenetic placement—nested within a large clade of exclusively African-endemic families rather than closely related to regionally sympatric North American or Neotropical lineages—was baffling. To test this counterintuitive phylogenetic and biogeographic hypothesis, we collected new specimens of *L. enigmatica* and sequenced, assembled, and annotated for the first time its complete mitochondrial genome. We then constructed the most taxonomically comprehensive mitochondrial data matrix of catfishes to date by including published mitochondrial genomes representing the majority of siluriform families. Using Bayesian co-estimation of phylogeny and divergence times, we inferred the evolutionary position of the puzzling *L. enigmatica* within a time-scaled global catfish phylogeny. Our results offer improved resolution and understanding of higher-level siluriform relationships and refine the timescale of catfish evolution. Crucially, our findings corroborate the hypothesis that *L. enigmatica* is sister to the African family Claroteidae and represents a relict lineage that originated in Africa during the Late Cretaceous (74.99 Ma; 95% HPD=68.58–81.39) and eventually culminated in Middle America. Our results therefore uphold the necessity of transoceanic dispersal—as opposed to Gondwanan vicariance—to explain this otherwise puzzling biogeographic pattern.

## 1. Introduction

Twenty years ago, an unusual and morphologically distinct catfish from Mexico’s Lacantún River (Usumacinta basin) was discovered and formally described, necessitating the erection of a new genus (*Lacantunia*) and family (Lacantuniidae), as it could not be assigned to any subordinal taxonomic ranks comprising the ∼3,000 recognized species of Siluriformes at the time (Rodiles-Hernández et al., 2005). The species epithet—*enigmatica*—reflects the puzzling taxonomic and evolutionary uncertainty posed by this catfish lineage, which baffled both its describers and the broader community of catfish systematists upon its discovery. The initial effort by the describing authors to resolve the phylogenetic placement of the Chiapas catfish, *Lacantunia enigmatica*, using comparative morphological data was inconclusive. Although this analysis excluded a close relationship between *L. enigmatica* and any sympatric catfish families (i.e., Ictaluridae, Heptapteridae, and Ariidae) (Rodiles-Hernández et al., 2005), the resulting unpublished phylogenetic assessment resolved *Lacantunia* crownward of diplomystids, hypsidorids and cetopsids, but unresolved with respect to other major monophyletic subgroups of siluriform families (Lundberg et al., 2007).

Shortly after the description of *L. enigmatica*, a molecular phylogenetic study aimed at shedding light on its phylogenetic placement and origins (Lundberg et al., 2007) used comparative molecular data consisting of two nuclear gene fragments (*rag1* and *rag2*, >3 kb total) sampled across more than 100 species representing most siluriform families, drawn from a then-contemporary study aimed at resolving higher-level relationships among catfishes globally (Sullivan et al., 2006). In addition to offering the first hypothesis for the phylogenetic placement of *Lacantunia* within the broader siluriform radiation, Lundberg et al. (2007) estimated divergence times across the resulting phylogeny—including the split between this enigmatic lineage and its closest living relative—using both Penalized Likelihood and Bayesian relaxed clock approaches as implemented in the software *r8s* (Sanderson, 2003) and *multidivtime* (Thorne and Kishino, 2002), respectively. The resulting phylogenetic and temporal inferences were perhaps as surprising and puzzling as the discovery of the Chiapas catfish itself, indicating that *Lacantunia* is nested within a multi-family clade of African freshwater catfishes and diverged from its closest living relative—the African-endemic family Claroteidae—sometime during the Late Cretaceous (∼85 Ma) (Lundberg et al., 2007). To account for this striking case of biogeographic disjunction, occurring too recently to be explained by Gondwanan vicariance, Lundberg et al. (2007) invoked dispersal via an ancient Holarctic intercontinental passage during the Late Cretaceous/Late Paleogene, followed by southward continental migration into Middle America.

Despite the existence of two subsequent molecular phylogenetic studies that have included sequence data from *L. enigmatica* (Chen et al., 2013; Peart et al., 2024), it could be argued that Lundberg et al. (2007) constitutes the only study explicitly addressing both the placement and timing of origin of the enigmatic Chiapas catfish within a comprehensive higher-level siluriform phylogeny. Nearly twenty years since its publication, testing the hypothesis of a Late Cretaceous origin and African roots of *Lacantunia* with independent comparative data is long overdue. To address this perceived deficiency, we first collected new specimens of *L. enigmatica*, and then sequenced, assembled, and annotated its first complete mitochondrial genome. By leveraging published complete mitochondrial genomes from representatives of most catfish families, we assembled the most taxonomically comprehensive molecular data matrix of catfishes to date. We then analyzed this matrix using phylogenetic inference and divergence time estimation methods to better understand the position, interrelationships, and evolutionary origin of *L. enigmatica*.

## 2. Methods

### 2.1. Specimen Sampling

The voucher specimen of *L. enigmatica* used to generate the complete mitochondrial genome for this study (Fig. 1), plus two additional individuals, were collected by JA, CL, and AAG at El Remolino (16°14’26” N, 90°50’53” W), a deep pool on the main channel of the Lacantún River (Usumacinta River basin) (Fig. 2). The site is located ∼3 km upriver from the small village of Reforma Agraria (Chiapas, Mexico), near the species’ type locality, and notable for the presence of whirlpools (hence its name in Spanish; remolino = whirlpool). Fishes were caught via deep angling using fresh shrimp as bait in the evening hours (∼9:00 p.m.) of 13 March 2025, the last night of a four-day dedicated collecting trip to the region. Specimens were euthanized with MS-222 in accordance with recommended guidelines for the use of fishes in research (Jenkins et al., 2014), prior to tissuing and preservation. Tissue samples for DNA extraction (fin clips) were taken prior to specimen fixation, preserved in 95% ethanol, flash frozen in liquid nitrogen while in the field, and subsequently cryopreserved long-term at -80 °C. Specimens were fixed in a 10% formalin solution and subsequently gradually transferred to 70% ethanol for long-term storage in the fish collections of Instituto de Biología, Universidad Nacional Autónoma de México (CNPE-IBUNAM) and El Colegio de la Frontera Sur, Unidad San Cristóbal de Las Casas (ECOSC). The specimen used in this study has been deposited at CNPE-IBUNAM under catalog number CNPE-IBUNAM 24453 (voucher JA2392). Specimens were collected under collecting permit request DGOPA-DAPA.-02764/25 addressed to the `Comisión Nacional de Acuacultura` (CONAPESCA).

**Figure 1.**
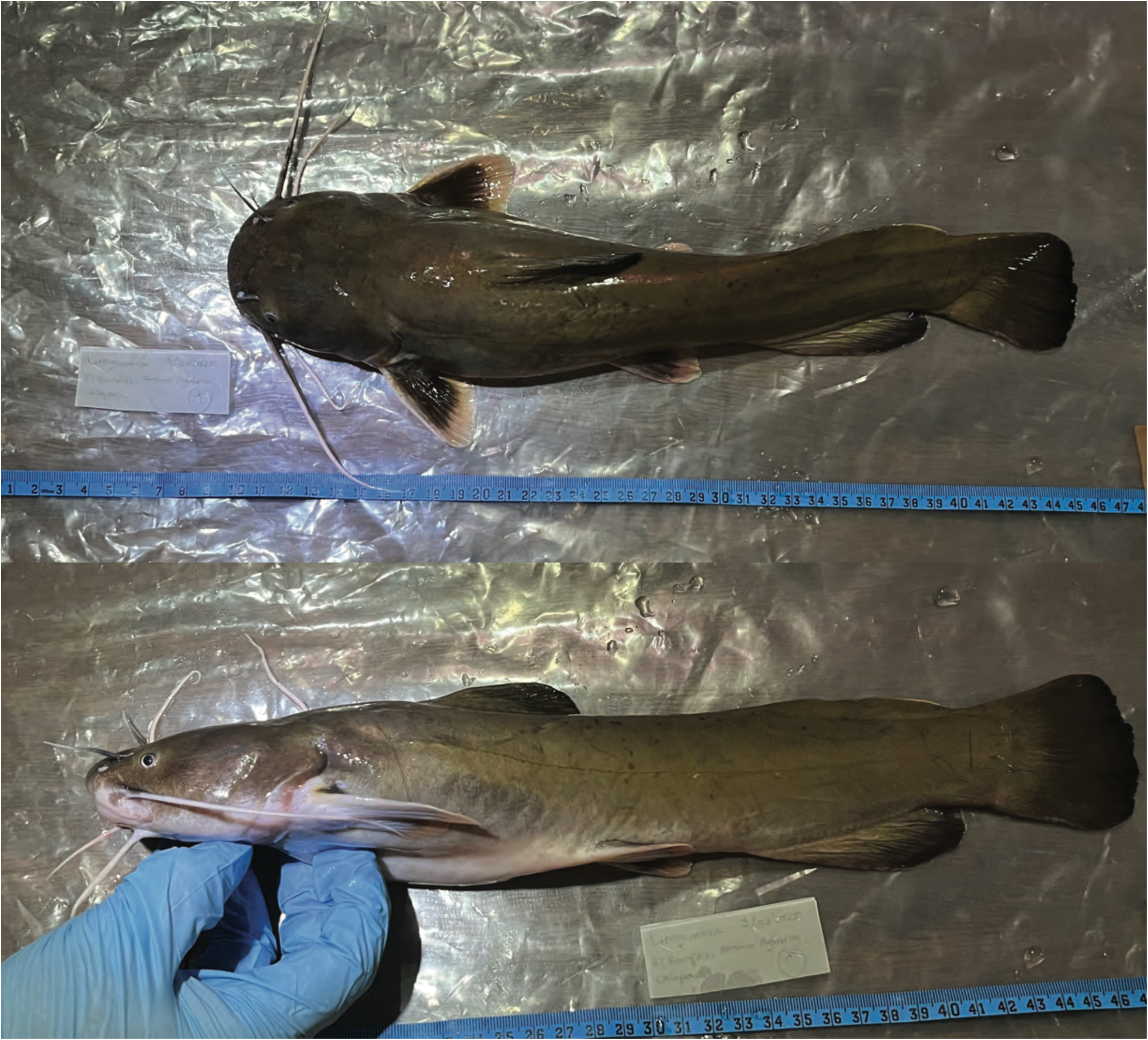
Voucher specimen. Live specimen of *Lacantunia enigmatica* (CNPE-IBUNAM 24453, voucher JA2392) in dorsal and lateral views. Photo taken in the field by JA and CLM immediately after collection.

**Figure 2.**
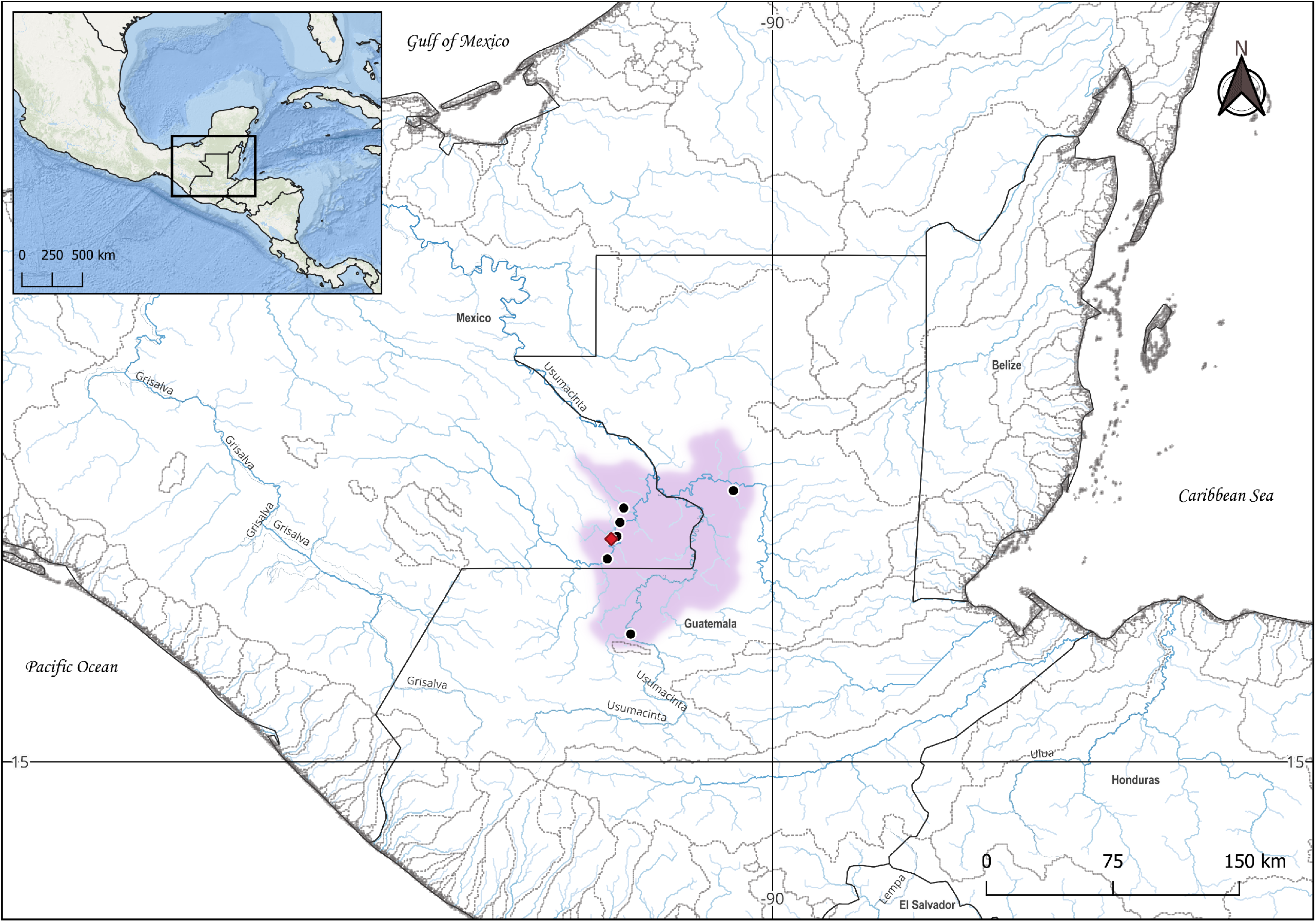
Study area. Map indicating the voucher’s sampling locality on the Lacantún River (red rhombus), known distribution records (black dots), and estimated geographic range (area in light purple) of *Lacantunia enigmatica* in the Middle Usumacinta River Basin. Rivers are shown in blue and basins are outlined with gray dotted boundaries.

### 2.2 Mitogenome sequencing, assembly, and annotation

Prior to library preparation for Illumina sequencing, total genomic DNA was extracted from the voucher’s cryopreserved tissue sample using a DNeasy Blood and Tissue Extraction Kit (Qiagen) following the manufacturer’s protocol. Extracted DNA was fragmented via sonication using a Diagenode Bioruptor® Pico device for eight cycles of alternating 20 s ultrasonic bursts and 30 s pauses in a 4°C water bath. Library preparation was carried out with a KAPA HyperPrep Kit (Kapa Biosystems) using 200 ng of fragmented DNA, quantified with an Invitrogen Qubit 4 Fluorometer. Fragmented DNA was end-repaired and end-polished via A-tailing and subsequently ligated to custom TruSeq dual-indexed adapters (Glenn et al., 2016) via short PCR. Products were size selected in a ∼300–500 bp range using magnetic beads, enriched through PCR (using the abovementioned kit), purified, and normalized. The resulting mitogenome library was sequenced in an Illumina HiSeq X at an average sequencing depth of 50X, producing 150-bp paired-end reads. Illumina sequencing was conducted at the Georgia Genomics and Bioinformatics Core of the University of Georgia (Athens, GA). Raw data quality was first evaluated in FastQC (Andrews, 2010), after which paired reads were trimmed to remove adapters using Geneious Prime v2025.2 (https://www.geneious.com). Mitogenome assembly was carried out using the module “map to reference” in Geneious, using three siluriform complete mitochondrial genomes available in GenBank as reference genomes for mapping reads during assembly: *Pangasius nasutus* OP236030 (Abdul Halim et al., 2023), *Chrysichthys nigrodigiatus* MH709123 (Kim et al., 2018), and *Rhamdia laticauda* PV662371 (Arroyave et al., 2025). These mitogenomes were used one at a time during assembly, thus generating three consensus mitogenomes which were then aligned in Mega X (Kumar et al., 2018) to generate a consensus mitogenome for *L. enigmatica*. Gene identification and annotation of the resulting mitochondrial genome was accomplished using MitoFish and MitoAnnotator v2025.06 (https://mitofish.aori.u-tokyo.ac.jp) (Iwasaki et al., 2013).

### 2.3 Taxon sampling, mitogenome mining, and matrix assembly

Ingroup taxa were sampled based on the availability of complete siluriform mitochondrial genomes in NCBI GenBank nucleotide database (https://www.ncbi.nlm.nih.gov/nucleotide/), with an emphasis on maximizing family-level representation. Similarly, outgroup taxa were sampled based on mitogenome availability while ensuring representation of all other otophysan orders (Characiformes, Cithariniformes, Gymnotiformes, and Cypriniformes) and from the order Gonorynchiformes, sister lineage of Otophysi, for rooting purposes. In total, 155 complete mitochondrial genomes from representatives of 34 of the 41 currently recognized living siluriform families (Van Der Laan et al., 2025) were downloaded from GenBank through a custom Python script. The complete mitochondrial genome of *L. enigmatica*, generated herein, was added to this set of ingroup mitogenomes, resulting in an ingroup consisting of 156 species from 35 catfish families (Supplementary Table 1). A total of 19 outgroup mitogenomes (from seven characiforms, five gymnotiforms, five cypriniforms, one cithariniform, and one gonorynchiform) were similarly downloaded from GenBank (Supplementary Table 1). Subsequently, the 13 protein-coding genes of each mitogenome were extracted using the metadata contained in each accession. Individual (by-gene) alignments were assembled with MAFFT v7.526 (Rozewicki et al., 2019) under the recommended settings for global alignments (--globalpair --maxiterate 1000), and subsequently concatenated with 2matrix v1.0 (Salinas and Little, 2014) into a data matrix consisting of 156 terminals and 12,994 aligned base pairs.

### 2.4 Bayesian co-estimation of phylogeny and divergence times

The phylogenetic placement and temporal context for the origin of *L. enigmatica* were investigated using a Bayesian relaxed phylogenetics approach. Prior to phylogeny inference, the best-fit model of molecular evolution for our global catfish mitogenome-wide data matrix (GTR+I+G) was determined with jModelTest v2.1.10 (Darriba et al., 2012) under the Akaike Information Criterion (AIC). Bayesian co-estimation of phylogeny and divergence times was conducted in BEAST2 v2.7.8 (Bouckaert et al., 2019) under a node-dating approach (Drummond et al., 2006), using an optimized relaxed clock (Douglas et al., 2020) and a birth–death (BD) speciation tree prior. The molecular clock was calibrated with fossil-based age estimates from 13 nodes across the ostariophysan clade—nine of which are within the Siluriformes—and incorporated using log-normally distributed priors (see Supplementary Methods 1 for a detailed account of the calibration strategy, including the list of calibration nodes, associated supporting paleontological evidence, proposed hard minimum and soft maximum age constraints, and justification for the temporal constraints and for the phylogenetic placement of calibration fossils).

The MCMC algorithm was run for 600 million generations and a sampling period of 1,000 generations. MCMC convergence was assessed via ESS values >200 using Tracer v1.7 (Rambaut et al., 2018), resulting in 25% of MCMC samples discarded as burn-in. The Maximum Clade Credibility (MCC) tree was obtained based on the set of post-burn-in samples using TreeAnnotator v2.7.7 (Bouckaert et al., 2019), resulting in a chronogram indicating posterior probabilities (PP) and mean ages of all nodes with their associated 95% highest posterior density (HPD) intervals. The topology was rooted at *Gonorynchus abbreviatus*.

## 3. Results

### 3.1 Mitogenome size and organization

The complete mitochondrial genome of *Lacantunia enigmatica* presented herein (GenBank accession PX522361) is 16,512 bp in total length, with an overall base composition biased toward A+T (A=32.6%, T= 28.0%, G= 13.6%, and C= 25.8%), and consisting of 37 genes of which 13 are protein coding, two are rRNAs, 22 are tRNAs, and one corresponds to the non-coding Control Region (CR)/D-loop (Fig. 3). The composition and arrangement of mitochondrial genes and associated features are detailed in Supplementary Table 2.

**Figure 3.**
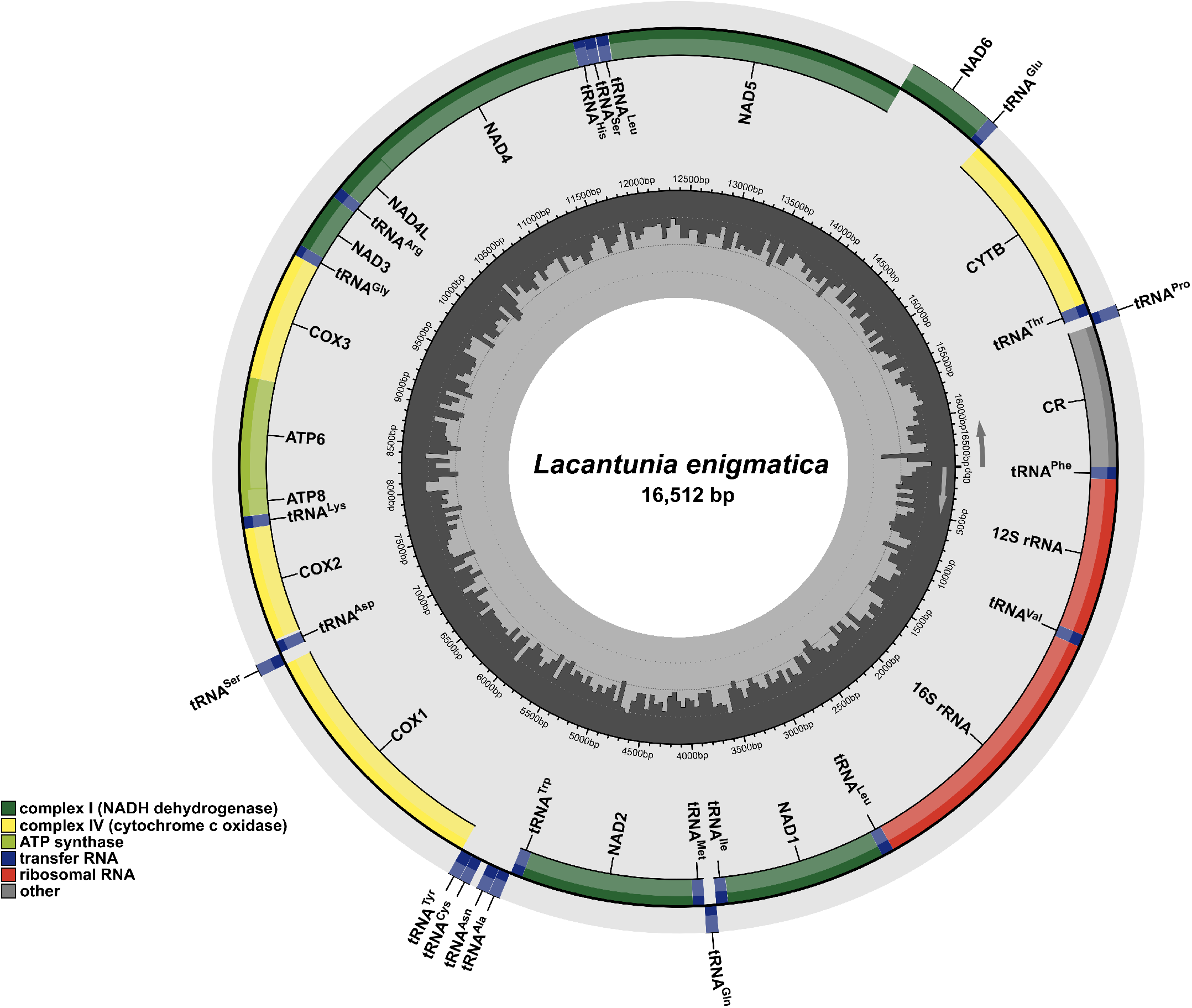
Mitochondrial genome map. Annotated map of the mitochondrial circular genome of *Lacantunia enigmatica* (GenBank accession PX522361). The outer ring corresponds to the L-(outermost) and H-strands, and depicts the location of protein-coding genes (all encoded in the H-strand, except for ND6), the non-coding control region (CR) (in dark grey), tRNAs (in blue), and rRNAs (in red). The inner ring (gray sliding window) denotes GC content along the genome.

### 3.2 Siluriform higher-level relationships and divergence times

Overall, the resulting global catfish time-scaled phylogeny (presented in full in Fig. 4 and as a simplified, family-level tree in Fig. 5) is robust, with most nodes strongly supported i.e., displaying posteriors probabilities (PP) greater than 0.98. The resulting topology splits Siluriformes into two reciprocally monophyletic lineages: one comprising the suborder Loricarioidei, and the other comprising the suborder Siluroidei and the family Diplomystidae, themselves resolved as sister taxa.

**Figure 4.**
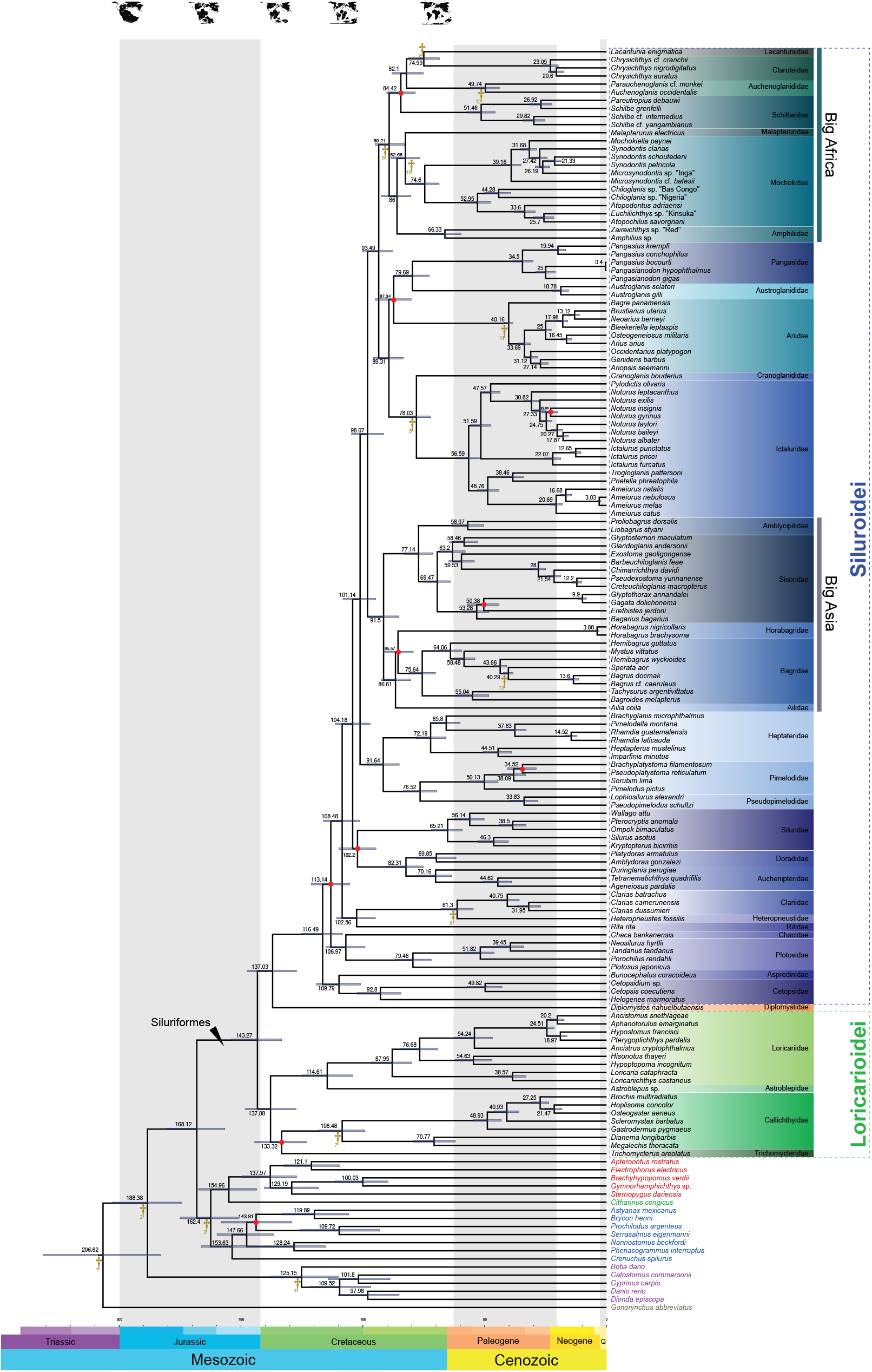
Time-scaled global phylogeny of Siluriformes. Chronogram resulting from Bayesian co-estimation of phylogeny and divergence times in the software BEAST based on mitogenomic comparative data from representatives of 35 of the 41 currently recognized living siluriform families. Fossil-based calibration nodes are indicated by gold crosses and follow the numbering presented in the calibration strategy (see Supplementary Methods 1). Most nodes are strongly supported i.e., with posterior probabilities (PP) larger than 0.98. Weakly supported nodes (PP≤0.98) are displayed in red. Divergence-time estimates are indicated by the mean ages of clades and the 95% highest posterior density (HPD) intervals (light blue bars) of mean node ages.

**Figure 5.**
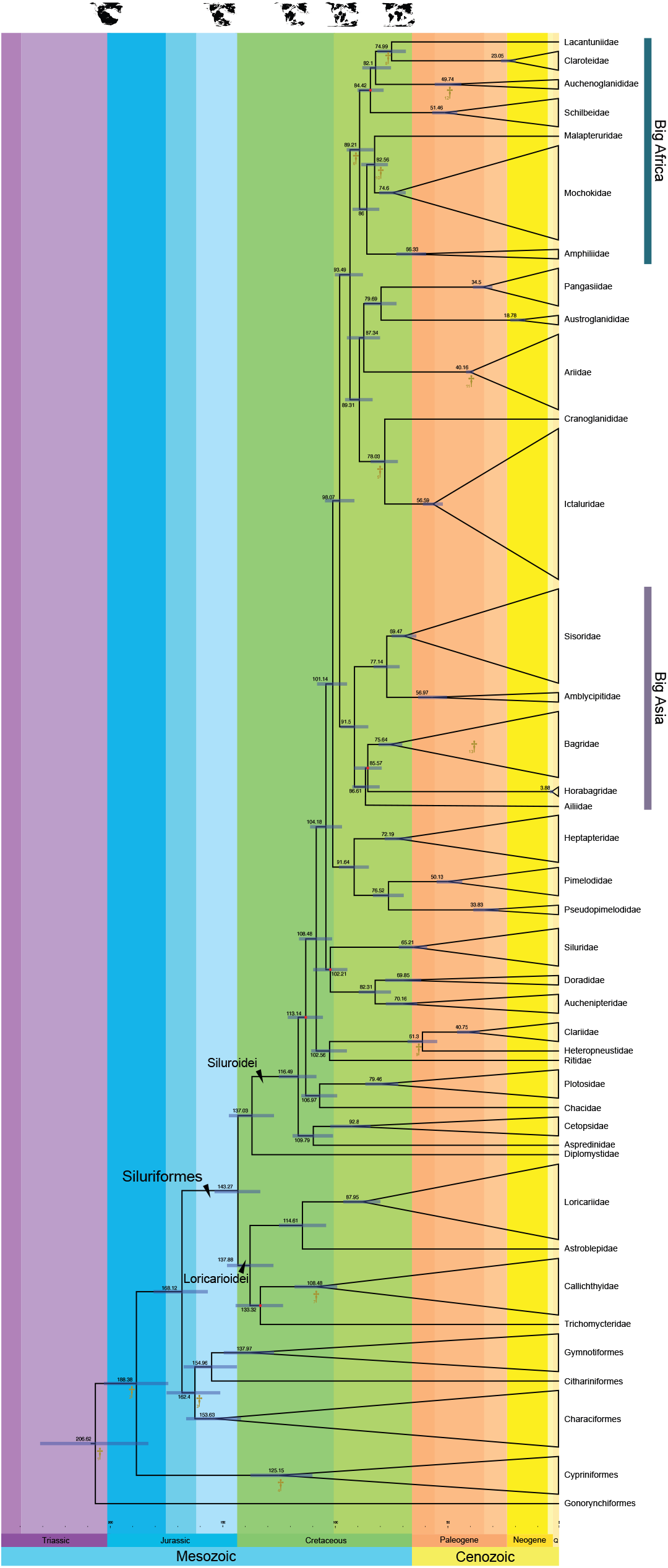
Time-scaled interfamilial siluriform relationships. Simplified representation of the chronogram shown in Figure 4, where major family-level clades are collapsed and depicted as triangles with widths proportional to the number of sampled terminals. The symbology follows that in Figure 4.

Our results support the monophyly of all sampled catfish families for which multiple samples were included. In contrast, the monophyly of genera where we sampled more than one species was only partially supported. Specifically, our results corroborated the monophyly of the genera *Chrysichthys, Chiloglanis, Austroglanis, Noturus, Ictalurus, Ameiurus, Horabagrus, Bagrus, Clarias*, and *Rhamdia*, whereas the genera *Synodontis, Microsynodontis, Schilbe, Pangasius*, and *Pangasiodon* were resolved as paraphyletic. The paraphyly of *Synodontis* and *Microsynodontis*, however, resulted from the placement of an undescribed taxon, *Microsynodontis* sp. “Inga”.

Within the family-rich Siluroidei, the large, continentally endemic multifamily clades informally known as “Big Africa” and “Big Asia”, were resolved with strong support (PP=1) and phylogenetically distant i.e., they belong to separate, deeply diverging lineages. The “Big Africa” clade was in turn resolved into two major subclades: one consisting of the families Malapteruridae, Mochokidae, and Amphiliidae, and another of Schilbeidae, Auchenoglanididae, Claroteidae, and Lacantuniidae. Notably, *Lacantunia enigmatica* was inferred as one of the main constituent lineages of the “Big Africa” clade, strongly supported as sister to Claroteidae (PP=1).

Regarding divergence time estimates, our resulting time-scaled phylogeny implies a Late Triassic origin for ostariophysan fishes (206.62 Ma; 95% HPD=183.06– 231.22) and a Late Jurassic/Early Cretaceous origin for Siluriformes (143.27 Ma; 95% HPD=133.32–153.74), with the suborders Siluroidei (116.49 Ma; 95% HPD=108.34– 125.15) and Loricarioidei (137.88 Ma; 95% HPD=127.44–148.16) both dating to the Early Cretaceous. Furthermore, our results entail that the “Big Africa” clade originated during the Late Cretaceous (89.21 Ma; 95% HPD=82.96–95), and the divergence between *Lacantunia* and its extant sister lineage (Claroteidae) also occurred in the Late Cretaceous approximately 15 Ma later (74.99 Ma; 95% HPD=68.58–81.39). Our results also indicate that most catfish family-level clades are of Paleogene origin or older, with some even extending to the Late Cretaceous (i.e., Mochokidae, Amphiliidae, Sisoridae, Bagridae, Heptapteridae, Doradidae, Auchenipteridae, Loricariidae, Cetopsidae, Callichthyidae).

## 4. Discussion

### 4.1 Catfish higher-level phylogenetics and interfamilial relationships

The breadth of our taxon and nucleotide sampling produced one of the most comprehensive siluriform phylogenies to date, providing new insights into higher-level relationships within catfishes. While our primary focus was on determining the phylogenetic position and divergence time of *Lacantunia enigmatica*, the resulting dataset and phylogeny allowed meaningful comparisons with previous hypotheses.

Siluriform higher-level interrelationships have been investigated based on broad taxonomic samplings since the early 1990s, initially using morphological data (Britto, 2003; de Pinna, 1993, 1998; Mo, 1991) and subsequently with DNA sequence data (Hardman, 2005, 2002; Kappas et al., 2016; Lundberg et al., 2007; Peart et al., 2024; Sullivan et al., 2006). Molecular phylogenetic studies in the mid-2000s corroborated several clades and relationships suggested by morphological data, while also uncovering novel relationships. Some of these studies also obtained molecular-clock-based estimates of absolute times of divergence, thus offering important insights into our understanding of catfish evolution. Most previous molecular studies, however, were based on Sanger-sequenced datasets and therefore relied on relatively short nucleotide alignments derived from a small number of loci. Among these, the study of Lundberg et al. (2007) is particularly relevant for comparison, as it is based on the same two-gene nuclear dataset of Sullivan et al. (2006) with the addition of nucleotide data from *Lacantunia enigmatica*.

Despite nearly two decades since the advent of high-throughput sequencing technologies (Bentley et al., 2008; Margulies et al., 2005) and their widespread application in higher-level fish systematics studies over the past decade (e.g., Arcila et al., 2017; Astudillo-Clavijo et al., 2023; Chakrabarty et al., 2017; Chen et al., 2012; Eytan et al., 2015; Faircloth et al., 2013; Harrington et al., 2024; Hughes et al., 2018; Melo et al., 2021; Melo and Stiassny, 2024), global catfish interrelationships have yet to be investigated under a phylogenomic framework. By leveraging complete mitochondrial genomes from representatives of most nominal living catfish families, our study provides what could be considered the most comprehensive phylogenomic assessment of catfish higher-level relationships. Notably, our data matrix represents nearly a fivefold increase in the number of nucleotides relative to the previously most taxonomically encompassing dataset (Lundberg et al., 2007). Increasing the amount of comparative nucleotide data has been shown to improve phylogenetic resolution and branch support in both theoretical (Martyn and Steel, 2012) and empirical studies (Kimball and Braun, 2014). Consistent with these expectations, our expanded dataset yielded higher overall increased phylogenetic resolution and more strongly supported clades (most nodes with PP≥0.98) with respect to previous studies (Hardman, 2005; Lundberg et al., 2007; Sullivan et al., 2006).

Our finding of Siluriformes consisting of three major clades, namely Siluroidei, Loricarioidei, and Diplomystidae, is consistent with previous studies. However, contrary to the prevailing morphology-based hypothesis positing the primitive Diplomystidae as the sister taxon to all other catfishes, our results support a sister-group relationship between Diplomystidae and Siluroidei. This finding corroborates the hypothesis first proposed by Sullivan et al. (2006) based on two nuclear loci, which initially surprised many catfish systematists (de Pinna et al., 2025). Although our corroboration of Sullivan et al.’s (2006) finding regarding the phylogenetic placement of Diplomystidae may suggest that this signal is rooted in molecular data, a more recent study using a 10-gene dataset, analyzed using methods to reduce evolutionary rate heterogeneity among lineages, supported a sister-group relationship between Diplomystidae and all other catfishes (Rivera-Rivera and Montoya-Burgos, 2018). This study attributed the discrepancy between morphological and previous molecular datasets to long branch attraction artifacts. Given that morphological evidence regarding the position of Diplomystidae remains somewhat conflicting (de Pinna et al., 2025), resolving the deepest nodes in the siluriform tree will likely remain challenging until whole-genome data provide new insights.

Regarding relationships within Loricarioidei, our results corroborate with strong support (PP=1) the well-established sister-group relationship between loricariids and astroblepids. They also suggest a relatively novel arrangement—albeit with moderate support (PP=0.94)—in which trichomycterids are sister to callichthyids. The latter finding, while not entirely in conflict with the conclusions of Sullivan et al. (2006)—in which trichomycterids are ambiguously resolved with respect to the remaining loricarioids (due to a polytomy)—is in disagreement with most morphology-based hypotheses and a recent synthesis and proposal of catfish higher-level relationships in which trichomycterids are the sister group of a clade consisting of all loricarioids except Nematogenyidae (de Pinna et al., 2025). Since our sampling lacked representatives from the families Nematogenyidae and Scoloplacidae, we cannot discount the possibility that our placing of Trichomycteridae may be influenced by our incomplete sampling of loricarioid major lineages.

A recent synthesis on the phylogenetic relationships of the major groups of Siluriformes (de Pinna et al., 2025) proposed that the suboder Siluroidei consists of eight superfamily-level clades, of which the Malapteruroidea and Bagroidea are roughly compositionally equivalent to the informally named “Big Africa” and “Big Asia” clades of Sullivan et al. (2006), respectively. As with Loricarioidei, within Siluroidei our results reveal instances of both agreement and conflict with the most relevant previous hypotheses, including the proposal of de Pinna et al. (2025). Previous studies have repeatedly suggested the family Cetopsidae is the sister group of the rest of Siluroidei (Britto, 2003; de Pinna, 1993, 1998; de Pinna et al., 2025; Lundberg et al., 2007), a view that is largely supported by our results. However, we identified and unexpected sister-group relationship (PP=0.99) between cetopsids and *Bunocephalus coracoideus*, the only sample from Aspredinidae included in our analysis. The prevailing view holds that Aspredinidae, together with the sister taxa Doradidae and Auchenipteridae, form a clade referred to as the superfamily Aspredinoidea (de Pinna et al., 2025; Sullivan et al., 2006). Given this context, our placement of *B. coracoideus* appears problematic. After cross-checking several genes of the mitogenome of *B. coracoideus* (AP012006) via NCBI’s BLAST, we could not confirm contamination or misidentification issues, discounting the possibility of a legacy error carried forward from the original source (Nakatani et al., 2011). A plausible explanation for this apparently spurious pattern is that it is an artifact of long-branch attraction. While further investigation of this conflicting result is beyond the scope of this study, our findings coupled with the fact that Aspredinoidea has a particularly convoluted history (de Pinna et al., 2025), underscore the need for improved hypothesis testing based on a larger taxon sampling within these families.

Although a close relationship between the families Plotosidae and Chacidae has been acknowledged since Bleeker’s times (Bleeker, 1863), it was not only until recently that they were explicitly proposed as sister taxa on the grounds of morphological evidence via cladistic analysis (de Pinna et al., 2025; Peixoto, 2018). Our study therefore constitutes the first DNA-based corroboration of this sister-group relationship. Our resultant phylogeny, however, failed to resolve these families forming a clade together with Siluridae and Clariidae, that is, the superfamily Siluroidea of de Pinna et al. (2025). Conversely, our results add to an ample body of studies independently supporting a clade consisting of the families Pimelodidae, Pseudopimelodidae, and Heptapteridae, which together with Phreatobiidae and *Conorhynchos* (not included here due to a lack of published mitogenomes) make up the superfamily Pimelodoidea (de Pinna et al., 2025; Sullivan et al., 2013).

The recent siluriform higher-level phylogenetic classification proposed by de Pinna et al. (2025) also includes two additional siluroid superfamily-level clades, namely Arioidea and Pangasioidea. Regarding the limits, composition, and placement of these two superfamilial clades, our results differ from the proposal of de Pinna et al. (2025) in two main ways. First, our phylogeny includes a well-supported clade (PP=1) consisting of all the families contained in Arioidea and Pangasioidea (except Anchariidae, for which there was no available mitogenome), thus negating the possibility of Arioidea being sister to Malapteruroidea, as proposed by de Pinna et al. (2025). Second, our inferred phylogenetic placement of Pangasiidae with respect to the remaining families in Arioidea and Pangasioidea goes against the definition of Pangasioidea. Specifically, our phylogeny resolves Pangasiidae as sister to Austroglanididae (PP=0.99), and together these families form a clade that is sister to Ariidae (albeit poorly supported; PP=0.53), a topology consistent with the results of a recent mitogenomic study focused on Pangasiidae (Duong et al., 2023). Conversely, in the proposal of de Pinna et al. (2025), Pangasiidae is sister to the clade Ictaluridae + Cranoglanididae, with all three families forming the superfamily Pangasioidea. By resolving Pangasiidae as the sister group of the “Big Africa” clade, the study of Lundberg et al. (2007) further contributes to the ambiguity regarding the position of Pangasiidae. In contrast to the uncertainty regarding the phylogenetic placement of Pangasiidae, the sister-group relationship between Ictaluridae and Cranoglanididae is a well-corroborated hypothesis (de Pinna et al., 2025; Lundberg et al., 2007; Sullivan et al., 2006) further supported by our findings (PP=1).

The existence of a siluroid superfamily-level clade consisting exclusively of Asian-endemic families (the “Big Asia” clade) was first proposed by Sullivan et al. (2006) and later largely corroborated by Kappas et al. (2016) and de Pinna et al. (2025), the latter who formally designated it as the superfamily Bagroidea. Our results strongly support the limits and composition of Bagroidea (PP=1) while offering similar yet slightly improved resolution with respect to the hypothesis of Sullivan et al. (2006) and the proposal of de Pinna et al. (2025). Likewise, our results offer strong support (PP=1) and corroboration for a large clade of mostly African-endemic families, which was first proposed by Sullivan et al. (2006) under the name “Big Africa” and corroborated by subsequent studies by Lundberg et al. (2007) and Peart et al. (2024) based on legacy markers and including *L. enigmatica*, and Kappas et al. (2016) and Schedel et al. (2022) also based on mitogenomes, but lacking data from *L. enigmatica*. This clade was further validated and designated as the superfamily Malapteruroidea by de Pinna et al. (2025). Crucially, by being based on a matrix consisting of mitogenome-scale data (∼13k aligned nucleotides) from a comprehensive and highly dense taxonomic sampling (156 species from 35 catfish families, including *L. enigmatica*), our study represents a significant improvement compared to these previous molecular studies.

Notably, the limits and interfamilial relationships within the “Big Africa” clade inferred with our mitogenomic dataset mirror the results of Lundberg et al. (2007) based on nDNA (genes *rag1* and *rag2*), including the puzzling inclusion of the Middle American *Lacantunia enigmatica* within this otherwise African-endemic clade (see subsection below). Similarly, except for the placement of Malapteruridae with respect to Amphiliidae and Mochokidae, our results match the findings of Peart et al. (2024) regarding the composition and resolution of this suprafamilial clade. While our findings agree with Schedel et al. (2022) on the constituent lineages of “Big Africa” (except for *L. enigmatica*, not included), their inferred intrafamilial relationships are not entirely congruent with ours or those reported in previous studies. By contrast, the limited taxonomic sampling of the “Big Africa” clade by Kappas et al. (2016) prevents meaningful comparison with our results.

Given conflicting results in key morphological studies regarding the phylogenetic placement of some families included in Malapteruroidea (de Pinna, 1993; Mo, 1991), our independent corroboration of the major findings from landmark molecular phylogenies (Lundberg et al., 2007; Peart et al., 2024; Sullivan et al., 2006) suggests that, in contrast to the extensive homoplasy displayed by morphological characters, comparative molecular data contain a strong signal for confidently resolving most major catfish lineages, including the “Big Africa” clade.

### 4.2 Phylogenetic origins of *Lacantunia* and the timeframe of catfish evolution

The initial assessment of the phylogenetic position and divergence timing of *L. enigmatica* (Lundberg et al., 2007) yielded indubitably unexpected results, warranting further verification to rule out potential methodological or analytical errors. To our knowledge, only two subsequent molecular phylogenetic studies (Chen et al., 2013; Peart et al., 2024) have included sequence data from *L. enigmatica*, making them the only empirical tests of Lundberg et al.’s (2007) findings to date. These studies used the same nuclear *rag1* and *rag2* sequence data from of Lundberg et al. (2007) with the addition of data for other taxa from nuclear markers *egr1, egr2b, egr3* and *rh1* (Chen et al., 2013) and from the mitochondrial *cytb* and *co1* and the nuclear *enc1* and *plagl2* (Peart et al., 2024). These studies, however, focused on ostariophysan higher-level relationships (Chen et al., 2013) and Lake Tanganyika catfish radiations (*Synodontis* and Claroteinae) (Peart et al., 2024), respectively, and therefore only tangentially addressed the phylogenetic position and origins of *Lacantunia*. Notably, their results neither fully replicate the findings of Lundberg et al. (2007) nor completely agree with each other. The limited sampling of Siluriformes in Chen et al. (2013) may reduce the confidence of their conclusions regarding *Lacantunia* and higher-level catfish relationships more broadly. Consequently, the study of Peart et al. (2024) provides the most rigorous test of Lundberg et al.’s (2007) findings prior to our study.

While Peart et al. (2024) corroborated the sister-group relationship between *Lacantunia* and Claroteidae, their divergence estimate for this split (54.32 Ma; 95% HPD=35.11–70.62) is younger than the Late Cretaceous age inferred by both Lundberg et al. (2007) (84.5 Ma; 95% HPD=75–94) and this study (74.99 Ma; 95% HPD=68.58– 81.39)—although the upper bound of Peart et al.’s HPD overlaps the Late Cretaceous. Notably, Peart et al. (2024) ascribed the discrepancy in age estimates to what they considered a suboptimal claroteid calibration in Lundberg et al. (2007). Specifically, in contrast to Lundberg et al. (2007), Peart et al. (2024) avoided using the claroteid calibration fossil †*Chrysichthys mahengeensis* as their analysis suggested that this fossil may not belong to the clade containing the type species of *Chrysichthys* and could not therefore be reliably placed on a phylogeny (Peart et al., 2014). Although our study did include †*Chrysichthys mahengeensis* among the 13 calibration fossils employed, our approach differed from Lundberg et al.’s (2007) in that we used this fossil to constrain the age of the MRCA of Claroteidae stem group instead of *Chrysichthys* stem group. This calibration decision was precisely based on the notion that generic attribution of †*Chrysichthys mahengeensis* remains ambiguous and requires further evidence (Otero, 2025; Peart et al., 2014) (see Supplementary Methods 1). It should also be noted that our 13-fossil calibration strategy included only three of the seven fossils used by Lundberg et al. (2007), supporting the argument that the calibrations are reasonably independent. As such, differences in calibration strategy, and/or data, likely contribute to the observed discrepancies in divergence estimates of this split across studies.

By resolving the Middle American *L. enigmatica* as the sister taxon of the family Claroteidae and deeply nested within an otherwise all-African suprafamilial clade, our results are noteworthy in that they provide independent empirical corroboration of this counterintuitive phylogenetic and biogeographic hypothesis. This confirmation strengthens confidence that previous hypotheses regarding the phylogenetic placement of *L. enigmatica* (Lundberg et al., 2007; Peart et al., 2024) reflect genuine signal. Beyond validating the African roots of *Lacantunia*, our study provides refined divergence-time estimates for the main constituent lineages of the catfish tree of life, including a Late Jurassic/Early Cretaceous age for the Siluriformes crown group (143.27 Ma; 95% HPD=133.32–153.74), Early Cretaceous origins for the suborders Siluroidei (116.49 Ma; 95% HPD=108.34–125.15) and Loricarioidei (137.88 Ma; 95% HPD=127.44–148.16), and Late Cretaceous ages for the “Big Africa” clade (89.21 Ma; 95% HPD=82.96–95) and the *Lacantunia*–Claroteidae split (74.99 Ma; 95% HPD=68.58–81.39). Notably, our estimate for the origin of the “Big Africa” clade closely matches that of Peart et al. (2024), who obtained 83.83 Ma (95% HPD = 66.76–93.16), demonstrating strong concordance at this deeper node. By contrast, the *Lacantunia*– Claroteidae split shows slightly greater variation among studies, reflecting the inherent uncertainty in dating this more recent divergence.

Overall, our divergence time estimates are largely consistent with those of Lundberg et al. (2007) and more recent molecular studies using mitogenomes (Kappas et al., 2016) and nuclear sequence data (Peart et al., 2024), providing a refined temporal context for catfish evolution. Despite this general concordance, however, our results imply a tendency toward older divergence times for the deepest nodes within Siluriformes, suggesting a more ancient origin for the major lineages than previously estimated. Interestingly, this pattern runs counter to the expectation that data sets with strong phylogenetic informativeness peaks near the present—such as protein-coding mitochondrial genes—tend to underestimate node ages at deeper divergences (Collins and Hrbek, 2018). Our slightly older estimates for the deepest nodes may therefore reflect the influence of our thorough fossil calibration strategy, particularly the inclusion of several well-supported deep-node constraints both inside and outside of catfishes (see Supplementary Methods 1). Importantly, by relying exclusively on fossil-based calibrations, our molecular clock analysis avoided propagating potentially inaccurate node ages from earlier molecular dating studies, thereby reducing bias in divergence-time estimation. Despite differences in data type (nuclear vs. mitochondrial DNA), calibration strategy, and analytical approaches, the general concordance in divergence-time estimates between our study and Lundberg et al.’s (2007) broadly substantiates the robustness of these temporal inferences. Indeed, the repeated recovery of broadly similar timelines across multiple datasets and methods suggests that the inferred evolutionary timescale is reliable (Duchêne et al., 2014; Mello et al., 2017).

### 4.3 Biogeographic considerations

Our time-scaled phylogeny reinforces the extraordinary case of biogeographic disjunction represented by the *Lacantunia*–Claroteidae split, here dated at ∼75 Ma (95% HPD=68.58–81.39) and therefore long postdating the Early Cretaceous separation of Africa and South America (McLoughlin, 2001). This timing supports the conclusion that divergence of the ancestral lineage leading to *L. enigmatica* cannot be explained by Gondwanan vicariance (Lundberg et al., 2007). Although the extant *L. enigmatica* is strictly freshwater, it remains possible that ancestral members of this lineage could tolerate brackish or marine environments (as most ariids and some plotosids and pangasiids can), which could have facilitated dispersal, with such tolerance subsequently lost, leaving *L. enigmatica* as the sole surviving freshwater representative. If the ancient lacantuniid lineage, however, was strictly freshwater, as are all extant members of the “Big Africa” clade, this would have rendered the possibility of direct marine dispersal unlikely. Under this scenario, explaining this otherwise puzzling biogeographic pattern requires invoking transoceanic dispersal via complex routes and mechanisms, including land bridges and/or stepping-stone pathways for continental dispersal and rafting for long-distance marine crossings (Capobianco and Friedman, 2019; Cavin, 2017). While individually improbable, such scenarios become increasingly plausible over deep evolutionary timescales and may help explain the remarkable intercontinental disjunction observed in the Claroteidae + *Lacantunia* clade. In fact, some of these dispersal mechanisms have been proposed to explain continentally disjunct distributions in other groups of Gondwanan freshwater fishes such as cichlids (Friedman et al., 2013; Matschiner, 2019; Matschiner et al., 2020, 2016), osteoglossids (Capobianco and Friedman, 2024), and cyprinodontiforms (Massip-Veloso et al., 2024). Critically, these taxa are examples of *secondary* freshwater fishes and therefore more tolerant to salinity than most catfish lineages (Cano-Barbacil et al., 2024).

To explain the biogeographic conundrum entailed by the inferred phylogenetic placement and divergence time of *Lacantunia*, Lundberg et al. (2007) hypothesized that the ancestral lineage leading to *L. enigmatica* dispersed from Africa to Middle America via an ancient Holarctic land bridge—either Beringia or the North Atlantic Land Bridge— aided by the warming and freshening of surface seawaters at critical passageways and times since the Late Cretaceous. This hypothetical biogeographic scenario, however, has yet to be supported by the discovery of either fossil or living intermediates in the lacantuniid lineage. While our findings uphold the necessity of transoceanic dispersal, the precise route, timing, and mechanism by which *L. enigmatica* reached Middle America from Africa remain as mysterious as the species’ epithet suggests.

## 5. Conclusions

Although the mitochondrial genome represents a single locus and can be susceptible to confounding effects of introgression, incomplete lineage sorting, and substitutional saturation, there is broad consensus that mitogenomic datasets have advanced the field of fish systematics (Miya and Nishida, 2015). Mitogenome-wide datasets have enabled the inference of robust phylogenetic hypotheses across multiple evolutionary timescales—from recent divergences to deep splits among major lineages—and have guided taxonomic and biogeographic inquiry in numerous fish groups (Aguilar et al., 2019; Arroyave et al., 2025; Maduna et al., 2022; Miya et al., 2003; Miya and Nishida, 2015; Santos et al., 2021; Schedel et al., 2022). The present study adds to this body of work, demonstrating the effectiveness of mitochondrial genomes for elucidating the evolutionary history of various fish clades. While complete resolution of deep divergences across the catfish tree remains challenging, our study offers improved resolution and support for key higher-level relationships and the timescale of siluriform evolution. Nevertheless, a canonical phylogenomic analysis—based on orders of magnitude more characters and loci—will likely be necessary to robustly resolve deep and contentious relationships resulting from short internodes and widespread morphological homoplasy.

Regarding the origins, phylogenetic affinities, and temporal context for the evolution of the sole extant representative of the family Lacantuniidae, *L. enigmatica*, our results strongly corroborate the findings of Lundberg et al. (2007) and Peart et al. 2024). Taken together, the available evidence indicates that the Chiapas catfish represents a relict lineage that originated in the Old World during the Late Cretaceous and subsequently underwent a complex yet poorly understood biogeographic history, culminating in its present restriction to a limited portion of the Middle Usumacinta River basin in the Guatemala-Mexico border region.

## 6. Acknowledgments

We would like to thank Dr. Eloisa Torres, Collection Manager at CNPE-IBUNAM, for her assistance with cataloging the voucher specimen of *Lacantunia enigmatica* used to generate the novel complete mitochondrial genome for this study.

## 8. Author contributions: CRediT

Conceptualization: JA, JJD, RAC Data curation: JA, AFMS, SHB, NRS

Formal analysis: JA, AFMS, SHB, NRS Funding acquisition: JA, JJD, AAGD

Investigation: JA, JJD, SHB, AFMS, CLM, AAGD, KK, RAC

Visualization: JA, AFMS, SHB, CLM

Project administration: JA, JJD

Writing – original draft: JA

Writing – review and editing: JA, JJD, RAC

## References

Abdul Halim, S.A.A., Esa, Y., Gan, H.M., Zainudin, A.A., Mohd Nor, S.A., 2023. The complete mitochondrial genomes of Pangasius nasutus and P. conchophilus (Siluriformes: Pangasiidae). Mitochondrial DNA B Resour. 8, 38–41. 10.1080/23802359.2022.2158694.

Aguilar, C., Miller, M.J., Loaiza, J.R., Krahe, R., De León, L.F., 2019. Mitogenomics of Central American weakly-electric fishes. Gene. 686, 164–170. 10.1016/j.gene.2018.11.045.

Andrews, S., 2010. FastQC: a quality control tool for high throughput sequence data. Babraham Bioinformatics, Babraham Institute, Cambridge, United Kingdom.

Arcila, D., Ortí, G., Vari, R., Armbruster, J.W., Stiassny, M.L.J., Ko, K.D., Sabaj, M.H., Lundberg, J., Revell, L.J., Betancur- R.R., 2017. Genome-wide interrogation advances resolution of recalcitrant groups in the tree of life. Nat. Ecol. Evol. 1, 0020. 10.1038/s41559-016-0020.

Arroyave, J., Mar-Silva, A.F., Melo, B.F., Hernández-Ávila, S.G., López-Vila, J.M., Silva, G.S.C., Díaz-Jáimes, P., 2025. Evolutionary history of Middle American Rhamdia (Siluriformes: Heptapteridae) inferred from comparative mitogenomic data: Insights on historical biogeography and cave colonization in the group. J. Syst. Evol. 63, 1501–1518. 10.1111/jse.70003.

Astudillo-Clavijo, V., Stiassny, M.L.J., Ilves, K.L., Musilova, Z., Salzburger, W., López-Fernández, H., 2023. Exon-based Phylogenomics and the Relationships of African Cichlid Fishes: Tackling the Challenges of Reconstructing Phylogenies with Repeated Rapid Radiations. Syst. Biol. 72, 134–149. 10.1093/sysbio/syac051.

Bentley, D.R., Balasubramanian, S., Swerdlow, H.P., Smith, G.P., Milton, J., Brown, C.G., Hall, K.P., Evers, D.J., Barnes, C.L., Bignell, H.R., Boutell, J.M., Bryant, J., Carter, R.J., Keira Cheetham, R., Cox, A.J., Ellis, D.J., Flatbush, M.R., Gormley, N.A., Humphray, S.J., Irving, L.J., Karbelashvili, M.S., Kirk, S.M., Li, H., Liu, X., Maisinger, K.S., Murray, L.J., Obradovic, B., Ost, T., Parkinson, M.L., Pratt, M.R., Rasolonjatovo, I.M.J., Reed, M.T., Rigatti, R., Rodighiero, C., Ross, M.T., Sabot, A., Sankar, S.V., Scally, A., Schroth, G.P., Smith, M.E., Smith, V.P., Spiridou, A., Torrance, P.E., Tzonev, S.S., Vermaas, E.H., Walter, K., Wu, X., Zhang, L., Alam, M.D., Anastasi, C., Aniebo, I.C., Bailey, D.M.D., Bancarz, I.R., Banerjee, S., Barbour, S.G., Baybayan, P.A., Benoit, V.A., Benson, K.F., Bevis, C., Black, P.J., Boodhun, A., Brennan, J.S., Bridgham, J.A., Brown, R.C., Brown, A.A., Buermann, D.H., Bundu, A.A., Burrows, J.C., Carter, N.P., Castillo, N., Chiara E. Catenazzi, M., Chang, S., Neil Cooley, R., Crake, N.R., Dada, O.O., Diakoumakos, K.D., Dominguez-Fernandez, B., Earnshaw, D.J., Egbujor, U.C., Elmore, D.W., Etchin, S.S., Ewan, M.R., Fedurco, M., Fraser, L.J., Fuentes Fajardo, K.V., Scott Furey, W., George, D., Gietzen, K.J., Goddard, C.P., Golda, G.S., Granieri, P.A., Green, D.E., Gustafson, D.L., Hansen, N.F., Harnish, K., Haudenschild, C.D., Heyer, N.I., Hims, M.M., Ho, J.T., Horgan, A.M., Hoschler, K., Hurwitz, S., Ivanov, D.V., Johnson, M.Q., James, T., Huw Jones, T.A., Kang, G.-D., Kerelska, T.H., Kersey, A.D., Khrebtukova, I., Kindwall, A.P., Kingsbury, Z., Kokko-Gonzales, P.I., Kumar, A., Laurent, M.A., Lawley, C.T., Lee, S.E., Lee, X., Liao, A.K., Loch, J.A., Lok, M., Luo, S., Mammen, R.M., Martin, J.W., McCauley, P.G., McNitt, P., Mehta, P., Moon, K.W., Mullens, J.W., Newington, T., Ning, Z., Ling Ng, B., Novo, S.M., O’Neill, M.J., Osborne, M.A., Osnowski, A., Ostadan, O., Paraschos, L.L., Pickering, L., Pike, Andrew C., Pike, Alger C., Chris Pinkard, D., Pliskin, D.P., Podhasky, J., Quijano, V.J., Raczy, C., Rae, V.H., Rawlings, S.R., Chiva Rodriguez, A., Roe, P.M., Rogers, John, Rogert Bacigalupo, M.C., Romanov, N., Romieu, A., Roth, R.K., Rourke, N.J., Ruediger, S.T., Rusman, E., Sanches-Kuiper, R.M., Schenker, M.R., Seoane, J.M., Shaw, R.J., Shiver, M.K., Short, S.W., Sizto, N.L., Sluis, J.P., Smith, M.A., Ernest Sohna, Sohna J., Spence, E.J., Stevens, K., Sutton, N., Szajkowski, L., Tregidgo, C.L., Turcatti, G., vandeVondele, S., Verhovsky, Y., Virk, S.M., Wakelin, S., Walcott, G.C., Wang, J., Worsley, G.J., Yan, J., Yau, L., Zuerlein, M., Rogers, Jane, Mullikin, J.C., Hurles, M.E., McCooke, N.J., West, J.S., Oaks, F.L., Lundberg, P.L., Klenerman, D., Durbin, R., Smith, A.J., 2008. Accurate whole human genome sequencing using reversible terminator chemistry. Nature. 456, 53–59. 10.1038/nature07517.

Bleeker, P., 1863. Systema Silurorum revisum. Ned. Tijdschr. Voor Dierkd. 1, 77–122.

Bouckaert, R., Vaughan, T.G., Barido-Sottani, J., Duchêne, S., Fourment, M., Gavryushkina, A., Heled, J., Jones, G., Kühnert, D., Maio, N.D., Matschiner, M., Mendes, F.K., Müller, N.F., Ogilvie, H.A., Plessis, L. du, Popinga, A., Rambaut, A., Rasmussen, D., Siveroni, I., Suchard, M.A., Wu, C.-H., Xie, D., Zhang, C., Stadler, T., Drummond, A.J., 2019. BEAST 2.5: An advanced software platform for Bayesian evolutionary analysis. PLoS Comput. Biol. 15, e1006650. 10.1371/journal.pcbi.1006650.

Britto, M.R. de, 2003. Análise filogenética da ordem siluriformes com ênfase nas relações da superfamília Loricarioidea (Teleostei: Ostariophysi). Universidade de Sao Paulo (USP).

Cano-Barbacil, C., Olden, J.D., García-Berthou, E., 2024. Myers’ divisions revisited: Contemporary evidence for distinct trait differences among global inland fishes. Fish Fish. 25, 672–685. 10.1111/faf.12832.

Capobianco, A., Friedman, M., 2024. Fossils indicate marine dispersal in osteoglossid fishes, a classic example of continental vicariance. Proc. Biol. Sci. 291, 20241293. 10.1098/rspb.2024.1293.

Capobianco, A., Friedman, M., 2019. Vicariance and dispersal in southern hemisphere freshwater fish clades: a palaeontological perspective. Biol. Rev. 94, 662–699. 10.1111/brv.12473.

Cavin, L., 2017. 150 million years of freshwater fish biogeography: vicariance or dispersal? Food Agric Sc Technol. 3, 1–4.

Chakrabarty, P., Faircloth, B.C., Alda, F., Ludt, W.B., Mcmahan, C.D., Near, T.J., Dornburg, A., Albert, J.S., Arroyave, J., Stiassny, M.L.J., Sorenson, L., Alfaro, M.E., 2017. Phylogenomic Systematics of Ostariophysan Fishes: Ultraconserved Elements Support the Surprising Non-Monophyly of Characiformes. Syst. Biol. 66, 881–895. 10.1093/sysbio/syx038.

Chen, M., Zou, M., Yang, L., He, S., 2012. Basal jawed vertebrate phylogenomics using transcriptomic data from Solexa sequencing. PloS One. 7, e36256. 10.1371/journal.pone.0036256.

Chen, W.-J., Lavoué, S., Mayden, R.L., 2013. Evolutionary origin and early biogeography of otophysan fishes (Ostariophysi: Teleostei). Evol. Int. J. Org. Evol. 67, 2218–2239. 10.1111/evo.12104.

Collins, R.A., Hrbek, T., 2018. An In Silico Comparison of Protocols for Dated Phylogenomics. Syst. Biol. 67, 633–650. 10.1093/sysbio/syx089.

Darriba, D., Taboada, G.L., Doallo, R., Posada, D., 2012. jModelTest 2: more models, new heuristics and parallel computing. Nat. Methods. 9, 772–772. 10.1038/nmeth.2109.

de Pinna, M.C.C., 1998. Phylogenetic Relationships of Neotropical Siluriformes □ : Historical Overview and Synthesis of Hypothesis, in: Malabarba, L.R., Reis, R.E., Vari, R.P., Lucena, C.A.S.de, Lucena, Z.de (Eds.), Phylogeny and Classification of Neotropical Fishes. EDIPUCRS, pp. 279–330.

de Pinna, M.C.C., 1993. Higher-level Phylogeny of Siluriformes, with a New Classification of the Order (Teleostei, Ostariophysi). The City University of New York.

de Pinna, M.C.C., Peixoto, L., Tagliacollo, V., Britto, M., 2025. Phylogenetic relationships and evolution of the major groups of Siluriformes, in: Arratia, G., Reis, R.E. (Eds), Catfishes, a Highly Diversified Group: Volume 2: Evolution and Phylogeny. CRC Press, pp. 97–127.

Douglas, J., Zhang, R., Bouckaert, R., 2020. Adaptive dating and fast proposals: revisiting the phylogenetic relaxed clock model. PloS Comput. Biol. 17, e1008322. 10.1101/2020.09.09.289124.

Drummond, A.J., Ho, S.Y.W., Phillips, M.J., Rambaut, A., 2006. Relaxed Phylogenetics and Dating with Confidence. PLoS Biol. 4, e88. 10.1371/journal.pbio.0040088.

Duchêne, S., Lanfear, R., Ho, S.Y.W., 2014. The impact of calibration and clock-model choice on molecular estimates of divergence times. Mol. Phylogenet. Evol. 78, 277–289. 10.1016/j.ympev.2014.05.032.

Duong, T.Y., Pham, L.T.K., Le, X.T.K., Nguyen, N.T.T., Nor, A.M., Le, T.H., 2023. Mitophylogeny of Pangasiid Catfishes and its Taxonomic Implications for Pangasiidae and the Suborder Siluroidei. Zool. Stud. 62, e48. 10.6620/ZS.2023.62-48.

Eytan, R.I., Evans, B.R., Dornburg, A., Lemmon, A.R., Lemmon, E.M., Wainwright, P.C., Near, T.J., 2015. Are 100 enough? Inferring acanthomorph teleost phylogeny using Anchored Hybrid Enrichment. BMC Evol. Biol. 15, 113. 10.1186/s12862-015-0415-0.

Faircloth, B.C., Sorenson, L., Santini, F., Alfaro, M.E., 2013. A Phylogenomic Perspective on the Radiation of Ray-Finned Fishes Based upon Targeted Sequencing of Ultraconserved Elements (UCEs). PLoS One. 8, e65923. 10.1371/journal.pone.0065923.

Friedman, M., Keck, B.P., Dornburg, A., Eytan, R.I., Martin, C.H., Hulsey, C.D., Wainwright, P.C., Near, T.J., 2013. Molecular and fossil evidence place the origin of cichlid fishes long after Gondwanan rifting. Proc. R. Soc. B Biol. Sci. 280, 20131733. 10.1098/rspb.2013.1733.

Glenn, T.C., Nilsen, R.A., Kieran, T.J., Finger, J.W., Pierson, T.W., Bentley, K.E., Hoffberg, S.L., Louha, S., León, F.J.G.-D., Portilla, M.A. del R., Reed, K.D., Anderson, J.L., Meece, J.K., Aggery, S.E., Rekaya, R., Alabady, M., Bélanger, M., Winker, K., Faircloth, B.C., 2016. Adapterama I: Universal Stubs and Primers for Thousands of Dual-Indexed Illumina Libraries (iTru & iNext). bioRxiv 049114. 10.1101/049114.

Hardman, M., 2005. The phylogenetic relationships among non-diplomystid catfishes as inferred from mitochondrial cytochrome b sequences; the search for the ictalurid sister taxon (Otophysi: Siluriformes). Mol. Phylogenet. Evol. 37, 700–720. 10.1016/j.ympev.2005.04.029.

Hardman, M., 2002. The phylogenetic relationships among extant catfishes, with special reference to Ictaluridae (Otophysi: Siluriformes). University of Illinois at Urbana-Champaign.

Harrington, R.C., Kolmann, M., Day, J.J., Faircloth, B.C., Friedman, M., Near, T.J., 2024. Dispersal sweepstakes: Biotic interchange propelled air-breathing fishes across the globe. J. Biogeogr. 51, 797–813. 10.1111/jbi.14781.

Hughes, L.C., Ortí, G., Huang, Y., Sun, Y., Baldwin, C.C., Thompson, A.W., Arcila, D., Betancur- R.R., Li, C., Becker, L., Bellora, N., Zhao, X., Li, X., Wang, M., Fang, C., Xie, B., Zhou, Z., Huang, H., Chen, S., Venkatesh, B., Shi, Q., 2018. Comprehensive phylogeny of ray-finned fishes (Actinopterygii) based on transcriptomic and genomic data. Proc. Natl. Acad. Sci. U.S.A. 115, 6249–6254. 10.1073/pnas.1719358115.

Iwasaki, W., Fukunaga, T., Isagozawa, R., Yamada, K., Maeda, Y., Satoh, T.P., Sado, T., Mabuchi, K., Takeshima, H., Miya, M., Nishida, M., 2013. MitoFish and MitoAnnotator: A Mitochondrial Genome Database of Fish with an Accurate and Automatic Annotation Pipeline. Mol. Biol. Evol. 30, 2531–2540. 10.1093/molbev/mst141.

Jenkins, J.A., Bart Jr, H.L., Bowker, J.D., Bowser, P.R., MacMillan, J.R., Nickum, J.G., Rose, J.D., Sorensen, P.W., Whitledge, G.W., Rachlin, J.W., 2014. Guidelines for the Use of Fishes in Research. Bethesda Md. USA Am. Fish. Soc.

Kappas, I., Vittas, S., Pantzartzi, C.N., Drosopoulou, E., Scouras, Z.G., 2016. A Time-Calibrated Mitogenome Phylogeny of Catfish (Teleostei: Siluriformes). PLoS One. 11, e0166988. 10.1371/journal.pone.0166988.

Kim, N.-K., Zealous Gietbong, F., Andriyono, S., Kim, A.R., Kim, H.-W., 2018. The complete mitogenome of Bagrid catfish Chrysichthys nigrodigitatus (Siluriformes: Claroteidae). Mitochondrial DNA B Resour. 3, 1239–1240. 10.1080/23802359.2018.1532341.

Kimball, R.T., Braun, E.L., 2014. Does more sequence data improve estimates of galliform phylogeny? Analyses of a rapid radiation using a complete data matrix. PeerJ. 2, e361. 10.7717/peerj.361.

Kumar, S., Stecher, G., Li, M., Knyaz, C., Tamura, K., 2018. MEGA X: Molecular Evolutionary Genetics Analysis across Computing Platforms. Mol. Biol. Evol. 35, 1547–1549. 10.1093/molbev/msy096.

Lundberg, J.G., Sullivan, J.P., Rodiles-Hernández, R., Hendrickson, D.A., 2007. Discovery of African roots for the Mesoamerican Chiapas catfish, Lacantunia enigmatica, requires an ancient intercontinental passage. Proc. Acad. Nat. Sci. Phila. 156, 39–53. 10.1635/0097-3157(2007)156%255B39:DOARFT%255D2.0.CO;2.

Maduna, S.N., Vivian-Smith, A., Jónsdóttir, Ó.D.B., Imsland, A.K.D., Klütsch, C.F.C., Nyman, T., Eiken, H.G., Hagen, S.B., 2022. Mitogenomics of the suborder Cottoidei (Teleostei: Perciformes): Improved assemblies, mitogenome features, phylogeny, and ecological implications. Genom. 114, 110297. 10.1016/j.ygeno.2022.110297.

Margulies, M., Egholm, M., Altman, W.E., Attiya, S., Bader, J.S., Bemben, L.A., Berka, J., Braverman, M.S., Chen, Y.-J., Chen, Z., Dewell, S.B., Du, L., Fierro, J.M., Gomes, X.V., Godwin, B.C., He, W., Helgesen, S., Ho, C.H., Irzyk, G.P., Jando, S.C., Alenquer, M.L.I., Jarvie, T.P., Jirage, K.B., Kim, J.-B., Knight, J.R., Lanza, J.R., Leamon, J.H., Lefkowitz, S.M., Lei, M., Li, J., Lohman, K.L., Lu, H., Makhijani, V.B., McDade, K.E., McKenna, M.P., Myers, E.W., Nickerson, E., Nobile, J.R., Plant, R., Puc, B.P., Ronan, .T., Roth, G.T., Sarkis, G.J., Simons, J.F., Simpson, J.W., Srinivasan, M., Tartaro, K.R., Tomasz, A., Vogt, K.A., Volkmer, G.A., Wang, S.H., Wang, Y., Weiner, M.P., Yu, P., Begley, R.F., Rothberg, J.M., 2005. Genome sequencing in microfabricated high-density picolitre reactors. Nature. 437, 376–380. 10.1038/nature03959.

Martyn, I., Steel, M., 2012. The impact and interplay of long and short branches on phylogenetic information content. J. Theor. Biol. 314, 157–163. 10.1016/j.jtbi.2012.08.040.

Massip-Veloso, Y., Hoagstrom, C.W., McMahan, C.D., Matamoros, W.A., 2024. Biogeography of Greater Antillean freshwater fishes, with a review of competing hypotheses. Biol. Rev. Camb. Philos. Soc. 99, 901–927. 10.1111/brv.13050.

Matschiner, M., 2019. Gondwanan vicariance or trans-Atlantic dispersal of cichlid fishes: a review of the molecular evidence. Hydrobiologia. 832, 9–37. 10.1007/s10750-018-3686-9.

Matschiner, M., Böhne, A., Ronco, F., Salzburger, W., 2020. The genomic timeline of cichlid fish diversification across continents. Nat. Commun. 11, 5895. 10.1038/s41467-020-17827-9.

Matschiner, M., Musilová, Z., Barth, J.M.I., Starostová, Z., Salzburger, W., Steel, M., Bouckaert, R., 2016. Bayesian Phylogenetic Estimation of Clade Ages Supports Trans-Atlantic Dispersal of Cichlid Fishes. Syst. Biol. syw076. 10.1093/sysbio/syw076.

McLoughlin, S., 2001. The breakup history of Gondwana and its impact on pre-Cenozoic floristic provincialism. Aust. J. Bot. 49, 271–300. 10.1071/bt00023.

Mello, B., Tao, Q., Tamura, K., Kumar, S., 2017. Fast and Accurate Estimates of Divergence Times from Big Data. Mol. Biol. Evol. 34, 45–50. 10.1093/molbev/msw247.

Melo, B.F., Sidlauskas, B.L., Near, T.J., Roxo, F.F., Ghezelayagh, A., Ochoa, L.E., Stiassny, M.L.J., Arroyave, J., Chang, J., Faircloth, B.C., MacGuigan, D.J., Harrington, R.C., Benine, R.C., Burns, M.D., Hoekzema, K., Sanches, N.C., Maldonado-Ocampo, J.A., Castro, R.M.C., Foresti, F., Alfaro, M.E., Oliveira, C., 2021. Accelerated Diversification Explains the Exceptional Species Richness of Tropical Characoid Fishes. Syst. Biol. 71, 78–92. 10.1093/sysbio/syab040.

Melo, B.F., Stiassny, M.L.J., 2024. Phylogenomic and anatomical evidence for the Late Cretaceous diversification of African characiform fishes, including a new family, under the influence of the Trans-Saharan Seaway. Evol. J. Linn. Soc. 3, kzae030. 10.1093/evolinnean/kzae030.

Miya, M., Nishida, M., 2015. The mitogenomic contributions to molecular phylogenetics and evolution of fishes: a 15-year retrospect. Ichthyol. Res. 62, 29–71. 10.1007/s10228-014-0440-9.

Miya, M., Takeshima, H., Endo, H., Ishiguro, N.B., Inoue, J.G., Mukai, T., Satoh, T.P., Yamaguchi, M., Kawaguchi, A., Mabuchi, K., Shirai, S.M., Nishida, M., 2003. Major patterns of higher teleostean phylogenies: a new perspective based on 100 complete mitochondrial DNA sequences. Mol. Phylogenet. Evol. 26, 121– 138. 10.1016/S1055-7903(02)00332-9.

Mo, T., 1991. Anatomy and systematics of Bagridae (Teleostei), and siluroid phylogeny. Koenigstein.

Nakatani, M., Miya, M., Mabuchi, K., Saitoh, K., Nishida, M., 2011. Evolutionary history of Otophysi (Teleostei), a major clade of the modern freshwater fishes: Pangaean origin and Mesozoic radiation. BMC Evol. Biol. 11, 177. 10.1186/1471-2148-11-177.

Otero, O., 2025. The African Fossil Catfishes, in: Arratia, G., Reis, R.E. (Eds), Catfishes, a Highly Diversified Group: Volume 2: Evolution and Phylogeny. CRC Press, pp. 34–53.

Peart, C.R., Bills, R., Newton, J., Near, T.J., Day, J.J., 2024. Do sympatric catfish radiations in Lake Tanganyika show eco-morphological diversification? Evol. J. Linn. Soc. 3, kzae015. 10.1093/evolinnean/kzae015.

Peart, C.R., Bills, R., Wilkinson, M., Day, J.J., 2014. Nocturnal claroteine catfishes reveal dual colonisation but a single radiation in Lake Tanganyika. Mol. Phylogenet. Evol. 73, 119–128. 10.1016/j.ympev.2014.01.013.

Peixoto, L.A.W., 2018. Implicações filogenéticas e taxonômicas na miologia facial comparada de Gymnotiformes e Siluriformes (Teleostei: Ostariophysi) (tese). Biblioteca Digital de Teses e Dissertações da Universidade de São Paulo. 10.11606/T.38.2018.tde-08052018-220803.

Rambaut, A., Drummond, A.J., Xie, D., Baele, G., Suchard, M.A., 2018. Posterior Summarization in Bayesian Phylogenetics Using Tracer 1.7. Syst. Biol. 67, 901– 904. 10.1093/sysbio/syy032.

Rivera-Rivera, C.J., Montoya-Burgos, J.I., 2018. Back to the roots: Reducing evolutionary rate heterogeneity among sequences gives support for the early morphological hypothesis of the root of Siluriformes (Teleostei: Ostariophysi). Mol. Phylogenet. Evol. 127, 272–279. 10.1016/j.ympev.2018.06.004.

Rodiles-Hernández, R., Hendrickson, D.A., Lundberg, J.G., Humphries, J.M., 2005. Lacantunia enigmatica (Teleostei: Siluriformes) a new and phylogenetically puzzling freshwater fish from Mesoamerica. Zootaxa. 1000, 1–24. 10.11646/zootaxa.1000.1.1.

Rozewicki, J., Li, S., Amada, K.M., Standley, D.M., Katoh, K., 2019. MAFFT-DASH: integrated protein sequence and structural alignment. Nucleic Acids Res. 47, W5–W10. 10.1093/nar/gkz342.

Salinas, N.R., Little, D.P., 2014. 2matrix: A utility for indel coding and phylogenetic matrix concatenation. Appl. Plant Sci. 2, 1300083. 10.3732/apps.1300083.

Sanderson, M.J., 2003. r8s: inferring absolute rates of molecular evolution and divergence times in the absence of a molecular clock. Bioinformatics. 19, 301– 302. 10.1093/bioinformatics/19.2.301.

Santos, R.P., Melo, B.F., Yazbeck, G.M., Oliveira, R.S., Hilário, H.O., Prosdocimi, F., Carvalho, D.C., 2021. Diversification of Prochilodus in the eastern Brazilian Shield: Evidence from complete mitochondrial genomes (Teleostei, Prochilodontidae). J. Zool. Syst. Evol. Res. 59, 1053–1063. 10.1111/jzs.12475.

Schedel, F.D.B., Chakona, A., Sidlauskas, B.L., Popoola, M.O., Usimesa Wingi, N., Neumann, D., Vreven, E.J.W.M.N., Schliewen, U.K., 2022. New phylogenetic insights into the African catfish families Mochokidae and Austroglanididae. J. Fish Biol. 100, 1171–1186. 10.1111/jfb.15014.

Sullivan, J.P., Lundberg, J.G., Hardman, M., 2006. A phylogenetic analysis of the major groups of catfishes (Teleostei: Siluriformes) using rag1 and rag2 nuclear gene sequences. Mol. Phylogenet. Evol. 41, 636–662. 10.1016/j.ympev.2006.05.044.

Sullivan, J.P., Muriel-Cunha, J., Lundberg, J.G., 2013. Phylogenetic Relationships and Molecular Dating of the Major Groups of Catfishes of the Neotropical Superfamily Pimelodoidea (Teleostei, Siluriformes). Proc. Acad. Nat. Sci. Phila. 162, 89–110. 10.1635/053.162.0106.

Thorne, J.L., Kishino, H., 2002. Divergence time and evolutionary rate estimation with multilocus data. Syst. Biol. 51, 689–702. 10.1080/10635150290102456.

Van Der Laan, R., Fricke, R., Eschmeyer, W.N., 2025. Eschmeyer’s catalog of fishes: classification [WWW Document]. URL https://www.calacademy.org/eschmeyers-catalog-of-fishes-classification (accessed 12.25.25).

