## Supplementary Methods 1 for "African origin and Late Cretaceous divergence of the Middle American catfish *Lacantunia enigmatica* corroborated by a global mitogenome phylogeny"

Below is the list of calibration nodes with their respective minimum age constraints, 95% soft maximum bounds, associated supporting paleontological evidence, and justifications for the temporal constraints and for the taxonomic/phylogenetic placement of calibration fossils. In all cases soft maximum ages were modeled by log-normally distributed priors.

**1. MRCA of pan-Ostariophysi.** Minimum age: 150 Ma. 95% soft maximum bound: 180 Ma. Calibration fossil: †*Tischlingerichthys viohli* (Arratia, 1997). Age and locality: Late Jurassic (150.94 Ma), Mörnsheim Formation of Mühlheim, Bavaria, Germany (Arratia, 1997; Benton et al., 2015). Phylogenetic and temporal justification: †*T. viohli* is widely regarded as the oldest pan-ostariophysan fossil and the sister taxon of the Ostariophysi crown group (Arratia, 1997, 2001; Benton et al., 2015; Near & Thacker, 2024). Although the origin and diversification of modern teleostean lineages has been long debated, a Middle-Late Jurassic minimum age for the Teleostei crown clade has been proposed based on paleontological evidence (Arratia, 1997, 2004), which, coupled with subsequent molecular dating studies that have recovered Middle Jurassic time estimates for the origin of modern ostariophysan fishes (Betancur-R. et al., 2013; Chen et al., 2013; Hughes et al., 2018; Near et al., 2013), support our proposed Early Jurassic soft maximum bound for the origin of the pan-ostariophysan lineage. Such an Early Jurassic maximum soft bound is further justified by the Toarcian Oceanic Anoxic Event, which resulted in the mass extinction of several early fish lineages but eventually promoted the post-extinction diversification of surviving ancestral lineages by filling vacated niches (Wignall, 2015).

**2. MRCA of pan-Otophysi.** Minimum age: 113 Ma. 95% soft maximum bound: 163 Ma. Calibration fossil: †*Santanichthys diasii* (da Silva Santos, 1958). Age and locality: Aptian and Albian, Early Cretaceous (100.5–121.4 Ma), Santana, Riachuelo, and Codó Formations, Romualdo Member, Brazil (da Silva Santos, 1958; Filleul & Maisey, 2004; Lindoso et al., 2016; R. M. Melo et al., 2020; Near & Thacker, 2024). Phylogenetic and temporal justification: †*S. diasii* was originally described as a leptolepid fish and subsequently redescribed as a stem characiform based on the presence of large and globular lagenar capsules, an autapomorphy of modern characiforms (Filleul & Maisey, 2004). However, in the same study, the authors also identified a series of plesiomorphic characters that reject characiform affinities for †*S. diasii*, including the presence of two supramaxillaries (absent in all living otophysans), a small supraoccipital crest (typical of early teleosts but well developed and posteriorly projected in characiforms), the absence of a dorsal depression in the quadrate (present in characiforms), the presence of free second ural and first preural centra (as in other stem otophysans), the fusion of the first hypural with the compound centrum (instead of the second hypural fused as in crown otophysans), and the absence of jaw teeth, which contrasts with the apomorphic multicuspid dentition of characiforms. A striking feature supporting the stem otophysan position of †*S. diasii* is the general architecture of the Weberian apparatus, which resembles that of other stem otophysans by lacking ossicles such as the claustrum, excluding the participation of the fifth vertebra, and retaining the plesiomorphic condition of the remaining bones (Diogo, 2009; Filleul & Maisey, 2004; Malabarba & Malabarba, 2010). The placement of †*S. diasii* as a stem otophysan is further supported by morphology-based phylogenetic analyses (Diogo, 2008; Mayrinck, 2011), leading to the recent recognition of this taxon as a pan-otophysan and the earliest fossil crown ostariophysan (Near & Thacker, 2024). Our proposed Late Jurassic 95% soft maximum bound for the origin of the pan-otophysan lineage is consistent with divergence time estimates from relevant time-scaled molecular phylogenies (Betancur-R. et al., 2013; Chen et al., 2013; Dai et al., 2018; Hughes et al., 2018; Near et al., 2012) as well as with the paleontological context. The Late Jurassic marks a major phase of diversification for early teleost fishes and a critical evolutionary turning point when teleosts began to radiate into many of their foundational lineages (Clarke & Friedman, 2018; Sferco et al., 2015). Consequently, it appears reasonable to probabilistically constrain the origin of otophysan fishes to the Late Jurassic.

**3. MRCA of pan-Characiformes.** Minimum age: 66 Ma. 95% soft maximum bound: 145 Ma. Calibration fossil: †*Tiupampichthys* *intermedius* (Gayet et al., 2003). Despite records of putative isolated characiform teeth from the Cenomanian of Sudan and Morocco (Malabarba & Malabarba, 2010), other fish lineages such as pycnodontids and lepisosteiforms may exhibit similar dentition and therefore the taxonomic assignation of these isolated teeth as characiform dentition is questionable (Cavin, 2017; Vullo et al., 2017). Consequently, the oldest confirmed characiform fossil consisting of both teeth and mandibular bones is that of †*T.* *intermedius*. Age and locality: Upper Maastrichtian, Late Cretaceous (66–70 Ma), El Molino Formation, Bolivia (Gayet et al., 2003). Phylogenetic and temporal justification: Recognition of †*T.* *intermedius* as a characiform is strongly supported by osteological characters (Gayet et al., 2003); however, its exact phylogenetic position within this clade remains unknown. Accordingly, the most conservative approach is to use this fossil to constrain the minimum age of the pan-Characiformes clade. Our proposed Early Cretaceous (145 Ma) 95% soft maximum bound is consistent with the results from a recent phylogenomic study that dated the Characiformes crown clade at 129.4 Ma (95% HPDI: 110–148.7 Ma) (B. F. Melo et al., 2021) and with the strongly supported biogeographic hypothesis that characiforms are Gondwanan in origin (Arroyave et al., 2013). Furthermore, if the Cenomanian fossil teeth were to be confirmed as characiform, this would extend the fossil record of the Characiformes to the Late Cretaceous, making an Early Cretaceous soft bound for the MRCA of pan-Characiformes quite reasonable.

**4. MRCA of crown Cypriniformes.** Minimum age: 60 Ma. 95% soft maximum bound: 140 Ma. Calibration fossil: Catostomidae *indet*. (left cleithrum), originally identified as Cyprinoidea *indet.* (Wilson, 1980) but subsequently assigned to the family Catostomidae (Liu, 2021; Liu et al., 2015). Age and locality: Tiffanian, Upper Paleocene (~58–60 Ma), Paskapoo Formation of Alberta, Canada (Wilson, 1980). Phylogenetic and temporal justification: Both the initial and subsequent taxonomic assignations concur that this this fossil correspond to a crown cypriniform, most likely with the family Catostomidae (Liu, 2021; Liu et al., 2015; Wilson, 1980). Our proposed 95% soft maximum bound (140 Ma) is consistent with most divergence time estimates for the origin of the Cypriniformes crown clade, according to which cypriniforms most likely originated at some point during the Early Cretaceous (Betancur-R. et al., 2013; Hughes et al., 2018; Near et al., 2012), while allowing the possibility—although with low probability—of a Late Jurassic origin, such as that proposed by Chen et al. (2013).

**5. MRCA of pan-Ictaluridae.** Minimum age: 59 Ma. 95% soft maximum bound: 95 Ma. Calibration fossil: †*Astephus* sp. (skull) (Grande & Lundberg, 1988; Lundberg, 1970, 1975, 1992). Age and locality: Tiffanian Ti3 North American Land Mammal Age (NALMA), Late Paleocene (59.5–60.3 Ma), Cedar Point Quarries of the Polecat Bench Formation, north-western Wyoming, USA (Higgins, 2003; Secord, 2008). Phylogenetic and temporal justification: Once held to be the sister taxon of extant Ictaluridae, and even assigned taxonomically to the family Ictaluridae (Lundberg, 1992), the position of †*Astephus* sp. relative to crown Ictaluridae has recently been revised, leading to its removal from Ictaluridae (Arce-H et al., 2017). Notwithstanding this phylogenetic reassessment, †*Astephus* sp. was inferred nested between crown Ictaluridae and Cranoglanididae (Arce-H et al., 2017), families that are recovered as sister in our topology. Consequently, †*Astephus* sp. was used herein to constrain the age of the split between Ictaluridae and Cranoglanididae, which in our topology is equivalent to the MRCA of pan-Ictaluridae. Our proposed soft maximum bound is compatible with the Late Cretaceous (~82 Ma) divergence between Ictaluridae and Cranoglanididae estimated by a previous molecular dating study (Lundberg et al., 2007).

**6. MRCA of the siluriform “Big Africa” clade.** Minimum age: 56 Ma. 95% soft maximum bound: 100 Ma. Calibration fossil: †*Nigerium wurnoense* (White, 1934). Age and locality: Thanetian, Late Paleocene (56–59 Ma), Dange Fomation, Wurno, Sokoto, Nigeria (Longbottom, 2010; Obaje, 2009; White, 1934). Phylogenetic and temporal justification: Although Longbottom (2010) placed †*Nigerium* in Claroteidae based on previous work by Mo (1991), the monophyly of Claroteidae *sensu* Mo was recently challenged by de Pinna et al. (2025). Given this emerging phylogenetic uncertainty, and following the recommendation of Otero (2025), †*Nigerium* *incerta sedis* is herein used to constrain the MRCA of the “Big Africa” clade *sensu* Sullivan et al. (2006). Our proposed soft maximum bound was informed by estimates for the age of this clade from arguably the most comprehensive global catfish time-scaled molecular phylogenies to date, which imply a Late Cretaceous (~80–95 Ma) origin for the “Big Africa” catfish lineage (Kappas et al., 2016; Lundberg et al., 2007).

**7. MRCA of crown Callichthyidae.** Minimum age: 56 Ma. 95% soft maximum bound: 80 Ma. Calibration fossil: †*Corydoras revelatus* (Cockerell, 1925). Age and locality: Thanetian, Late Paleocene (~56–58 Ma), Maiz Gordo Formation, Argentina (Campo et al., 2007; Cockerell, 1925). Phylogenetic and temporal justification: Despite being described in the genus *Corydoras*, the original generic attribution of †*C. revelatus* is not without caveats and has recently been refuted—on the basis of having a short mesethmoid (vs. long in *Corydoras*)—and tentatively attributed to *Hoplisoma* (Britto et al., 2025; Dias et al., 2025). Notwithstanding this taxonomic uncertainty, because it is the oldest known fossil unambiguously attributed to modern Callichthyidae, it was herein used to constrain the age of the callichthyid crown clade. Molecular clock estimates for the age of this clade are somewhat contrasting, with mean values that are either rather young (~60–70 Ma) (Lundberg et al., 2007; Mariguela et al., 2013) or fairly old (104 Ma) (Marburger et al., 2018). Our proposed Late Cretaceous soft maximum bound therefore offers a reasonable compromise between a K-Pg boundary and a mid-Cretaceous origin for this clade.

**8. MRCA of stem Claroteidae.** Minimum age: 45 Ma. 95% soft maximum bound: 85 Ma. Calibration fossil: †*Chrysichthys mahengeensis*. (Murray & Budney, 2003). Age and locality: Lutenian, Eocene (~45 Ma), Mahenge site, Tanzania (Murray & Budney, 2003; Uhl et al., 2024). Phylogenetic and temporal justification: While family-level assignation is unquestionable and supported by abundant anatomical evidence, attribution to the extant genus *Chrysichthys* was made to emphasize the resemblance to present-day members and is not supported by any apomorphy (Otero, 2025). Given that generic attribution requires further evidence, and that our sampling of claroteids consists only of species of *Chrysichthys*, †*C. mahengeensis* was herein used to constrain the age of the MRCA of Claroteidae stem group (as opposed to crown group). Our Late Cretaceous soft maximum bound is grounded in the findings of two benchmark studies on global siluriform phylogenetics, which agree on an Eocene (~53 Ma) origin for the Claroteidae *crown* group (Kappas et al., 2016; Lundberg et al., 2007).

**9. MRCA of stem Clariidae.** Minimum age: 45 Ma. 95% soft maximum bound: 66 Ma. Calibration fossils: Clariidae *incertae sedis* (pectoral spines, right anguloarticular) (Gayet et al., 1987). Age and locality: Early-Middle Eocene (45–50 Ma), Kuldana Formation, Kohat District, Pakistan (Gingerich, 2003). Phylogenetic and temporal justification: While the original taxonomic designation of the Kuldana fossil material as Clariidae *indet*. is unequivocal and supported by several anatomical diagnostic features related to the shape of the fossil bones (Gayet et al., 1987), an accurate phylogenetic placement within Clariidae of this material has never been proposed (Agnese & Teugels, 2005), most likely due to its fragmentary nature. Given this phylogenetic uncertainty, these Early-Middle Eocene fossil remains are herein used to constrain the age of the Clariidae stem clade. The divergence between Clariidae and Heteropneustidae (effectively the MRCA of stem Clariidae) has been dated by previous studies to the Paleogene, with mean estimates from the Eocene (~43 Ma) (Kappas et al., 2016) and from the Paleocene (~60 Ma) (Lundberg et al., 2007). Our proposed soft lower bound accounts for these previous estimates while allowing the possibility (although small) of an earlier (Late Cretaceous) origin. Notably, this soft maximum constraint also accounts for a fast recovery of freshwater systems after the Cretaceous-Paleogene extinction (Brownstein & Lyson, 2022; Robertson et al., 2013), which may have promoted the diversification of clariids and other contemporary fish lineages.

**10. MRCA of stem Mochokidae.** Minimum age: 38 Ma. 95% soft maximum bound: 90 Ma. Calibration fossils: Mochokidae *indet*. (isolated teeth) (Murray et al., 2010; Otero et al., 2015). Age and locality: Upper Bartonian, Middle Eocene (38–39 Ma), Dur At-Talah, Lybia (Jaeger et al., 2010; Otero et al., 2015); Priabonian, Late Eocene (37 Ma), Birket Qarun Formation, Locality 2 (BQ-2), Fayum Depression, Egypt (Murray et al., 2010; Seiffert et al., 2008). Phylogenetic and temporal justification: While their distinctive S shape with pointed tips allows for the unambiguous assignation of these fossil teeth to the family Mochokidae, it is not clear if they belong to a living genus (Murray et al., 2010; Otero et al., 2015). Accordingly, these fossil teeth were used herein to constrain the age of the Mochokidae stem group. Molecular clock-based estimates for the age of Mochokidae crown clade converge on a Late Cretaceous origin, near the K-Pg boundary (Day et al., 2023; Lundberg et al., 2007). Expectedly, the proposed divergence between Mochokidae and Malapteruridae (effectively the MRCA of stem Mochokidae) is older, estimated at ~85 Ma (Lundberg et al., 2007). Our suggested soft maximum bound therefore attempts to accommodate these divergence time estimates while minimizing an already long fossil gap.

**11. MRCA of crown Ariidae.** Minimum age: 37 Ma. 95% soft maximum bound: 66 Ma. Calibration fossil: †*Qarmoutus hitanensis* (El-Sayed et al., 2017). Despite reports of considerably older putative ariid fossils (Late Campanian/Early Maastrichtian) (Cione, 1987), their highly fragmentary and incomplete nature (mostly otoliths) casts doubt on their unequivocal taxonomic assignment (Betancur-R, 2009; Cione & Prasad, 2002). Meanwhile, †*Q. hitanensis* was described based on the oldest best preserved and most complete skeletal specimen reliably assignable to Ariidae, consisting of a partial neurocranium (the complete left side), partial right dentary, left suspensorium, two opercles, left pectoral girdle and spine, nuchal plates, first and second dorsal spines, Weberian apparatus, and a disassociated series of abdominal vertebrae (El-Sayed et al., 2017). Age and locality: Priabonian, Upper Eocene (~37 Ma), Birket Qarun Formation, Wadi El-Hitan, Fayum Depression, northern Western Desert, Egypt (El-Sayed et al., 2017). Phylogenetic and temporal justification: To our knowledge, there is no fossil specimen older than †*Q. hitanensis* that matches its level of completeness and preservation and is unmistakably attributable to Ariidae. Despite some conflicting hypotheses, most phylogenetic reconstructions place *Q. hitanensis* nested within Ariidae crown group (El-Sayed et al., 2017) and therefore it was used herein to constrain this clade. While Lundberg et al. (2007) inferred an Eocene (~40 Ma) mean age for the origin of Ariidae, Betancur (2009) proposed considerably older estimates (~57–88 Ma). These earlier divergence times, however, are likely strongly influenced by Betancur’s usage of the abovementioned Late Cretaceous problematic fossils. Therefore, our proposed soft maximum bound at the K-Pg boundary is conservative and more aligned with unambiguous fossil evidence—precluding a large fossil gap—while at the same time allowing the possibility of a Late Cretaceous origin for the Ariidae, although with low probability.

**12. MRCA of crown Auchenoglanididae.** Minimum age: 37 Ma. 95% soft maximum bound: 66 Ma. Calibration fossil: †*Auchenoglanis* sp. (two proximal portions of pectoral spines) (El-Sayed et al., 2020). Age and locality: Priabonian, Late Eocene (37 Ma), Birket Qarun Formation, Locality 2 (BQ-2), Fayum Depression, Egypt (El-Sayed et al., 2020; Murray et al., 2010; Sayed et al., 2025; Seiffert et al., 2008). Phylogenetic and temporal justification: Although these disarticulated fossil remains have been unambiguously attributed to the genus *Auchenoglanis* on account of their shape and ornamentation (El-Sayed et al., 2020; Otero, 2025), due to our data matrix including only one species of *Auchenoglanis*, these fossils were used herein to constrain the age of the MRCA of Auchenoglanidae as delimited by our taxon sampling (*Auchenoglanis* *occidentalis* + *Parauchenoglanis* cf. *monkei*. Our proposed K-Pg boundary soft maximum bound for Auchenoglanididae is consistent with the inferred Middle Eocene mean age estimate of Lundberg et al. (2007) while allowing a reasonable fossil gap (<30 Ma) and the possibility—although low—of a Late Cretaceous origin.

**13. MRCA of stem *Bagrus*.** Minimum age: 37 Ma. 95% soft maximum bound: 56 Ma. Calibration fossil: †*Bagrus* sp. (El-Sayed et al., 2020). Age and locality: Priabonian, Late Eocene (37 Ma), Birket Qarun Formation, Locality 2 (BQ-2), Fayum Depression, Egypt (El-Sayed et al., 2020; Murray et al., 2010; Sayed et al., 2025; Seiffert et al., 2008). Phylogenetic and temporal justification: Anatomical evidence strongly supports the placement of this fossil in the genus *Bagrus* (El-Sayed et al., 2020) but its exact phylogenetic position remains unclear. Therefore, †*Bagrus* sp. is used herein to constrain the stem *Bagrus* clade. Lundberg et al. (2007) inferred an Eocene (~40 Ma) mean divergence time between *Bagrus* and *Hemibagrus*, however, without consideration of the Eocene *Bagrus* fossil herein used for calibration purposes, unavailable at the time. To allow for the possibility of an older origin (Paleocene) for *Bagrus* while constraining the fossil gap, we used a soft maximum bound of 56 Ma.

**References**

Agnese, J.-F., Teugels, G.G., 2005. Insight into the phylogeny of African Clariidae (Teleostei, Siluriformes): Implications for their body shape evolution, biogeography, and taxonomy. Mol. Phylogenet. Evol. 36, 546–553. https://doi.org/10.1016/j.ympev.2005.03.028.

Arce-H, M., Lundberg, J.G., O’Leary, M.A., 2017. Phylogeny of the North American catfish family Ictaluridae (Teleostei: Siluriformes) combining morphology, genes and fossils. Cladistics 33, 406–428. https://doi.org/10.1111/cla.12175.

Arratia, G., 1997. Basal teleosts and teleostean phylogeny. Palaeo. Ichthyol. 7, 1–168.

Arratia, G., 2001. The Sister-Group of Teleostei: Consensus and Disagreements. J. Vertebr. Paleontol. 21, 767–773.

Arratia, G., 2004. Mesozoic halecostomes and the early radiation of teleosts, in: Arraita, G., Tintori, A. (Eds), Mesozoic fishes 3. Dr. Friedrich Pfeil, Munich, pp. 279–315.

Arroyave, J., Denton, J.S.S., Stiassny, M.L.J., 2013. Are characiform Fishes Gondwanan in Origin? Insights from a Time-Scaled Molecular Phylogeny of the Citharinoidei (Ostariophysi: Characiformes). PLoS One. 8, e77269. https://doi.org/10.1371/journal.pone.0077269.

Benton, M., Donoghue, P., Vinther, J., Asher, R., Friedman, M., Near, T., 2015. Constraints on the timescale of animal evolutionary history. Palaeontol. Electron. https://doi.org/10.26879/424.

Betancur-R, R., 2009. Molecular phylogenetics and evolutionary history of ariid catfishes revisited: a comprehensive sampling. BMC Evol. Biol. 9, 175. https://doi.org/10.1186/1471-2148-9-175.

Betancur-R., R., Broughton, R.E., Wiley, E.O., Carpenter, K., López, J.A., Li, C., Holcroft, N.I., Arcila, D., Sanciangco, M., Cureton II, J.C., Zhang, F., Buser, T., Campbell, M.A., Ballesteros, J.A., Roa-Varon, A., Willis, S., Borden, W.C., Rowley, T., Reneau, P.C., Hough, D.J., Lu, G., Grande, T., Arratia, G., Ortí, G., 2013. The Tree of Life and a New Classification of Bony Fishes. PLoS Curr. 5:ecurrents.tol.53ba26640df0ccaee75bb165c8c26288. https://doi.org/10.1371/currents.tol.53ba26640df0ccaee75bb165c8c26288.

Britto, M.R., Tencatt, L.F.C., Dopazo, M., Santos, S.A. dos, Ferreira, K.C.F. (in memoriam), Reis, R.E., 2025. Phylogeny and Classification of the Family Callichthyidae (Armored Catfishes), in: Arraita, G., Reis, R.E., (Eds), Catfishes, a Highly Diversified Group. CRC Press.

Brownstein, C.D., Lyson, T.R., 2022. Giant gar from directly above the Cretaceous–Palaeogene boundary suggests healthy freshwater ecosystems existed within thousands of years of the asteroid impact. Biol. Lett. 18, 20220118. https://doi.org/10.1098/rsbl.2022.0118.

Campo, M. do, Papa, C. del, Jiménez-Millán, J., Nieto, F., 2007. Clay mineral assemblages and analcime formation in a Palaeogene fluvial–lacustrine sequence (Maíz Gordo Formation Palaeogen) from northwestern Argentina. Sediment. Geol. 201, 56-74. https://doi.org/10.1016/j.sedgeo.2007.04.007

Cavin, L., 2017. Freshwater Fishes: 250 Million Years of Evolutionary History. Elsevier.

Chen, W.-J., Lavoué, S., Mayden, R.L., 2013. Evolutionary origin and early biogeography of otophysan fishes (Ostariophysi: Teleostei). Evol. 67, 2218–2239. https://doi.org/10.1111/evo.12104

Cione, A.L., 1987. The Late Cretaceous fauna of los Alamitos, Patagonia, Argentina. II: The fishes - La faune crétacé supérieur de Los Alamitos, Patagonie, Argentine. II. Les poissons. Revista del Museo Argentino de Ciencias Naturales Bernardino Rivadavia e Instituto Nacional de Investigacion de las Ciencias Naturales. Paleontologia 3, 111–120.

Cione, A.L., Prasad, G.V.R., 2002. The Oldest Known Catfish (Teleostei:Siluriformes) from Asia (India, Late Cretaceous). J. of Paleontol. 76, 190–193.

Clarke, J.T., Friedman, M., 2018. Body-shape diversity in Triassic–Early Cretaceous neopterygian fishes: sustained holostean disparity and predominantly gradual increases in teleost phenotypic variety. Paleobiology. 44, 402–433. https://doi.org/10.1017/pab.2018.8.

Cockerell, T.D.A., 1925. A Fossil Fish of the Family Callichthyidae. Science. 62, 397–398. https://doi.org/10.1126/science.62.1609.397.c.

da Silva Santos, R., 1958. *Leptolepis diasii*, novo peixe fóssil da Serra do Araripe, Brasil. Ministério da Agricultura, Departamento Nacional da Produção Mineral.

Dai, W., Zou, M., Yang, L., Du, K., Chen, W., Shen, Y., Mayden, R.L., He, S., 2018. Phylogenomic Perspective on the Relationships and Evolutionary History of the Major Otocephalan Lineages. Sci. Rep. 8, 205. https://doi.org/10.1038/s41598-017-18432-5.

Day, J.J., Steell, E.M., Vigliotta, T.R., Withey, L.A., Bills, R., Friel, J.P., Genner, M.J., Stiassny, M.L.J., 2023. Exceptional levels of species discovery ameliorate inferences of the biogeography and diversification of an Afrotropical catfish family. Mol. Phylogenet. and Evol. 182, 107754. https://doi.org/10.1016/j.ympev.2023.107754.

de Pinna, M.C.C., Peixoto, L., Tagliacollo, V., Britto, M., 2025. Phylogenetic relationships and evolution of the major groups of Siluriformes, in: Arraita, G., Reis, R.E., (Eds), Catfishes, a Highly Diversified Group. CRC Press.

Dias, A.C., Tencatt, L.F.C., Roxo, F.F., Silva, G. de S. da C., Santos, S.A., Britto, M.R., Taylor, M.I., Oliveira, C., 2025. Phylogenomic analyses in the complex Neotropical subfamily Corydoradinae (Siluriformes: Callichthyidae) with a new classification based on morphological and molecular data. Zool. J. Linn. Soc. 203, zlae053. https://doi.org/10.1093/zoolinnean/zlae053.

Diogo, R., 2009. Origin, Evolution and Homologies of the Weberian Apparatus: A New Insight. Int. J. of Morphol. 27, 333-354. https://doi.org/10.4067/S0717-95022009000200008.

Diogo, R., 2008. The Origin of Higher Clades: Osteology, Myology, Phylogeny and Evolution of Bony Fishes and the Rise of Tetrapods. CRC Press, Boca Raton. https://doi.org/10.1201/9780429063978.

El-Sayed, S., Kora, M.A., Sallam, H.M., Claeson, K.M., Seiffert, E.R., Antar, M.S., 2017. A new genus and species of marine catfishes (Siluriformes; Ariidae) from the upper Eocene Birket Qarun Formation, Wadi El-Hitan, Egypt. PLoS One, 12, e0172409. https://doi.org/10.1371/journal.pone.0172409.

El-Sayed, S., Murray, A.M., Kora, M.A., Abu El-Kheir, G.A., Antar, M.S., Seiffert, E.R., Sallam, H.M., 2020. Oldest Record of African Bagridae and Evidence from Catfishes for a Marine Influence in the Late Eocene Birket Qarun Locality 2 (BQ-2), Fayum Depression, Egypt. J. Vertebr. Paleontol. 40, e1780248. https://doi.org/10.1080/02724634.2020.1780248.

Filleul, A., Maisey, J.G., 2004. Redescription of *Santanichthys diasii* (Otophysi, Characiformes) from the Albian of the Santana Formation and comments on its implications for otophysan relationships. Am. Mus. Novit. 3455.

Gayet, M., De Broin, F., Rage, J.-C., 1987. Lower Vertebrates from the Early-Middle Eocene Kuldana Formation of Kohat (Pakistan): Holostei and Teleostei, Chelonia, and Squamata. Contrib. Mus. Pal. Univ. Mich. 27, 151–168.

Gayet, M., Jégu, M., Bocquentin, J., Negri, F.R., 2003. New characoids from the Upper Cretaceous and Paleocene of Bolivia and the Mio-Pliocene of Brazil: phylogenetic position and paleobiogeographic implications. J. Vertebr. Paleontol. 23, 28–46. https://doi.org/10.1671/0272-4634(2003)23[28:NCFTUC]2.0.CO;2.

Gingerich, P.D., 2003. Stratigraphic and micropaleontological constraints on the middle Eocene age of the mammal-bearing Kuldana Formation of Pakistan. J. Vertebr Paleontol. 23, 643–651. https://doi.org/10.1671/2409.

Grande, L., Lundberg, J.G., 1988. Revision and redescription of the genus *Astephus* (Siluriformes: Ictaluridae) with a discussion of its phylogenetic relationships. J. Vertebr. Paleontol. 8, 139–171. https://doi.org/10.1080/02724634.1988.10011694.

Higgins, P., 2003. A Wyoming succession of Paleocene mammal-bearing localities bracketing the boundary between the Torrejonian and Tiffanian North American Land Mammal “Ages.” Rocky Mountain Geology 38, 247–280. https://doi.org/10.2113/gsrocky.38.2.247.

Hughes, L.C., Ortí, G., Huang, Y., Sun, Y., Baldwin, C.C., Thompson, A.W., Arcila, D., Betancur-R., R., Li, C., Becker, L., Bellora, N., Zhao, X., Li, X., Wang, M., Fang, C., Xie, B., Zhou, Z., Huang, H., Chen, S., Venkatesh, B., Shi, Q., 2018. Comprehensive phylogeny of ray-finned fishes (Actinopterygii) based on transcriptomic and genomic data. Proc. Natl. Acad. Sci. U.S.A. 115, 6249–6254. https://doi.org/10.1073/pnas.1719358115.

Jaeger, J.-J., Marivaux, L., Salem, M., Bilal, A.A., Benammi, M., Chaimanee, Y., Duringer, P., Marandat, B., Métais, E., Schuster, M., Valentin, X., Brunet, M., 2010. New rodent assemblages from the Eocene Dur At-Talah escarpment (Sahara of central Libya): systematic, biochronological, and palaeobiogeographical implications. Zool. J. Linn. Soc. 160, 195–213. https://doi.org/10.1111/j.1096-3642.2009.00600.x.

Kappas, I., Vittas, S., Pantzartzi, C.N., Drosopoulou, E., Scouras, Z.G., 2016. A Time-Calibrated Mitogenome Phylogeny of Catfish (Teleostei: Siluriformes). PLoS One 11, e0166988. https://doi.org/10.1371/journal.pone.0166988.

Lindoso, R.M., Maisey, J.G., Carvalho, I. de S., 2016. Ichthyofauna from the Codó Formation, Lower Cretaceous (Aptian, Parnaíba Basin), Northeastern Brazil and their paleobiogeographical and paleoecological significance. Palaeogeogr. Palaeoclimatol. Palaeoecol. 447, 53–64. https://doi.org/10.1016/j.palaeo.2016.01.045.

Liu, J., 2021. Redescription of ‘Amyzon’ brevipinne and remarks on North American Eocene catostomids (Cypriniformes: Catostomidae). J. Sys Palaeontol. 19, 677–689. https://doi.org/10.1080/14772019.2021.1968966.

Liu, J., Chang, M.-M., Wilson, M.V.H., Murray, A.M., 2015. A new family of Cypriniformes (Teleostei, Ostariophysi) based on a redescription of †Jianghanichthys hubeiensis (Lei, 1977) from the Eocene Yangxi Formation of China. J. Vertebr. Paleontol. 35, e1004073. https://doi.org/10.1080/02724634.2015.1004073.

Longbottom, A., 2010. A new species of the catfish *Nigerium* from the Palaeogene of the Tilemsi Valley, Republic of Mali. Palaeontology. 53, 571–594. https://doi.org/10.1111/j.1475-4983.2010.00946.x.

Lundberg, J.G., 1992. The phylogeny of ictalurid catfishes: a synthesis of recent work. Systematics, historical ecology, and North American freshwater fishes. 392, 420.

Lundberg, J.G., 1975. The Fossil Catfishes of North America. Papers on paleontology.

Lundberg, J.G., 1970. The evolutionary history of North American catfishes, family Ictaluridae. University of Michigan.

Lundberg, J.G., Sullivan, J.P., Rodiles-Hernández, R., Hendrickson, D.A., 2007. Discovery of African roots for the Mesoamerican Chiapas catfish, *Lacantunia enigmatica*, requires an ancient intercontinental passage. Proc. Acad. Nat. Sci. Philadelphia. 156, 39–53.

Malabarba, L., Malabarba, M., 2010. Biogeography of Characiformes: an evaluation of the available information of fossil and extant taxa, in: Origin and Phylogenetic Interrelationships of Teleosts. Verlag Friedrich Pfeil, München, Germany, pp. 317–336.

Marburger, S., Alexandrou, M.A., Taggart, J.B., Creer, S., Carvalho, G., Oliveira, C., Taylor, M.I., 2018. Whole genome duplication and transposable element proliferation drive genome expansion in Corydoradinae catfishes. Proc. Biol. Sci. 285, 20172732. https://doi.org/10.1098/rspb.2017.2732.

Mariguela, T.C., Alexandrou, M.A., Foresti, F., Oliveira, C., 2013. Historical biogeography and cryptic diversity in the Callichthyinae (Siluriformes, Callichthyidae). J. Zool. Syst. Evol. Res. 51, 308–315. https://doi.org/10.1111/jzs.12029.

Mayrinck, D. de, 2011. Phylogenetic relationships of otophysan (Actinopterygii, Teleostei), notably the Characiformes, including fossil members. Cybium. 35, 74–75.

Melo, B.F., Sidlauskas, B.L., Near, T.J., Roxo, F.F., Ghezelayagh, A., Ochoa, L.E., Stiassny, M.L.J., Arroyave, J., Chang, J., Faircloth, B.C., MacGuigan, D.J., Harrington, R.C., Benine, R.C., Burns, M.D., Hoekzema, K., Sanches, N.C., Maldonado-Ocampo, J.A., Castro, R.M.C., Foresti, F., Alfaro, M.E., Oliveira, C., 2021. Accelerated Diversification Explains the Exceptional Species Richness of Tropical Characoid Fishes. Syst. Biol. 71, 78-92. https://doi.org/10.1093/sysbio/syab040.

Melo, R.M., Guzmán, J., Almeida-Lima, D., Piovesan, E.K., Neumann, V.H. de M.L., Sousa, A. de J. e, 2020. New marine data and age accuracy of the Romualdo Formation, Araripe Basin, Brazil. Sci. Rep. 10, 15779. https://doi.org/10.1038/s41598-020-72789-8

Mo, T., 1991. Anatomy and systematics of Bagridae (Teleostei), and siluroid phylogeny. Koenigstein.

Murray, A.M., Budney, L.A., 2003. A new species of catfish (Claroteidae, *Chrysichthys*) from an Eocene crater lake in East Africa. Can. J. Earth Sci. 40, 983–993. https://doi.org/10.1139/e03-031.

Murray, A.M., Cook, T.D., Attia, Y.S., Chatrath, P., Simons, E.L., 2010. A freshwater ichthyofauna from the late Eocene Birket Qarun Formation, Fayum, Egypt. J. Vertebr. Paleontol. 30, 665–680. https://doi.org/10.1080/02724631003758060

Near, T.J., Dornburg, A., Eytan, R.I., Keck, B.P., Smith, W.L., Kuhn, K.L., Moore, J.A., Price, S.A., Burbrink, F.T., Friedman, M., Wainwright, P.C., 2013. Phylogeny and tempo of diversification in the superradiation of spiny-rayed fishes. Proc. Natl. Acad. Sci. U.S.A. 110, 12738–12743. https://doi.org/10.1073/pnas.1304661110.

Near, T.J., Eytan, R.I., Dornburg, A., Kuhn, K.L., Moore, J.A., Davis, M.P., Wainwright, P.C., Friedman, M., Smith, W.L., 2012. Resolution of ray-finned fish phylogeny and timing of diversification. Proc. Natl. Acad. Sci. U.S.A. 109, 13698–13703. https://doi.org/10.1073/pnas.1206625109.

Near, T.J., Thacker, C.E., 2024. Phylogenetic Classification of Living and Fossil Ray-Finned Fishes (Actinopterygii). Bull. Peabody. Mus. Nat. Hist. 65, 3–302. https://doi.org/10.3374/014.065.0101.

Obaje, N.G., 2009. The Sokoto Basin (Nigerian Sector of the Iullemmeden Basin), in: Obaje, N.G. (Ed.), Geology and Mineral Resources of Nigeria. Springer, Berlin, Heidelberg, pp. 77–89. https://doi.org/10.1007/978-3-540-92685-6_7.

Otero, O., 2025. The African Fossil Catfishes, in: Catfishes, a Highly Diversified Group: Volume 2: Evolution and Phylogeny. CRC Press.

Otero, O., Pinton, A., Cappetta, H., Adnet, S., Valentin, X., Salem, M., Jaeger, J.-J., 2015. A Fish Assemblage from the Middle Eocene from Libya (Dur At-Talah) and the Earliest Record of Modern African Fish Genera. PLoS One. 10, e0144358. https://doi.org/10.1371/journal.pone.0144358.

Robertson, D.S., Lewis, W.M., Sheehan, P.M., Toon, O.B., 2013. K-Pg extinction patterns in marine and freshwater environments: The impact winter model. J. Geophys. Res. Biogeosci. 118, 1006–1014. https://doi.org/10.1002/jgrg.20086.

Sayed, M.M., Heinz, P., Abd El-Gaied, I.M., El-Kahawy, R.M., Sayed, D.M., Salama, Y.F., Al-Hashim, M.H., Wagreich, M., 2025. Paleobiodiversity, Paleobiogeography, and Paleoenvironments of the Middle–Upper Eocene Benthic Foraminifera in the Fayum Area, Western Desert, Egypt. J. Mar. Sci. Eng. 13, 663. https://doi.org/10.3390/jmse13040663.

Secord, R., 2008. The Tiffanian Land-Mammal Age (Middle and Late Paleocene) in the Northern Bighorn Basin, Wyoming. Department of Earth and Atmospheric Sciences: Faculty Publications.

Seiffert, E.R., Bown, T.M., Clyde, W.C., Simons, E., 2008. Geology, Paleoenvironment, and Age of Birket Qarun Locality 2 (BQ-2), Fayum Depression, Egypt, in: Fleagle, J.G., Gilbert, C.C. (Eds.), Elwyn Simons: A Search for Origins, Developments in Primatology: Progress and Prospects. Springer New York, New York, NY, pp. 71–86. https://doi.org/10.1007/978-0-387-73896-3_8.

Sferco, E., López-Arbarello, A., Báez, A.M., 2015. Phylogenetic relationships of ^†^*Luisiella feruglioi* (Bordas) and the recognition of a new clade of freshwater teleosts from the Jurassic of Gondwana. BMC Evol. Biol. 15, 268. https://doi.org/10.1186/s12862-015-0551-6.

Sullivan, J.P., Lundberg, J.G., Hardman, M., 2006. A phylogenetic analysis of the major groups of catfishes (Teleostei: Siluriformes) using rag1 and rag2 nuclear gene sequences. Mol. Phylogenet. Evol. 41, 636–662. https://doi.org/10.1016/j.ympev.2006.05.044.

Uhl, D., Wuttke, M., Aiglstorfer, M., Gee, C.T., Grandi, F., Höltke, O., Kaiser, T.M., Kaulfuss, U., Lee, D., Lehmann, T., Oms, O., Poschmann, M.J., Rasser, M.W., Schindler, T., Smith, K.T., Suhr, P., Wappler, T., Wedmann, S., 2024. Deep-time maar lakes and other volcanogenic lakes as Fossil-Lagerstätten – An overview. Palaeobiodivers. Palaeoenviron. 104, 763–848. https://doi.org/10.1007/s12549-024-00635-0.

Vullo, R., Cavin, L., Khalloufi, B., Amaghzaz, M., Bardet, N., Jalil, N.-E., Jourani, E., Khaldoune, F., Gheerbrant, E., 2017. A unique Cretaceous–Paleogene lineage of piranha-jawed pycnodont fishes. Sci. Rep. 7, 6802. https://doi.org/10.1038/s41598-017-06792-x.

White, E.I., 1934. Fossil fishes of Sokoto province. Published by the Authority of the Nigerian Government, [Lagos].

Wignall, P.B., 2015. The Worst of Times : How Life on Earth Survived Eighty Million Years of Extinctions 1–224.

Wilson, M.V.H., 1980. Oldest known *Esox* (Pisces: Esocidae), part of a new Paleocene teleost fauna from western Canada. Can. J. Earth Sci. 17, 307–312. https://doi.org/10.1139/e80-030.
