## Supplementary Table 1 for "African origin and Late Cretaceous divergence of the Middle American catfish *Lacantunia enigmatica* corroborated by a global mitogenome phylogeny"

**Supplementary Table 1.** List of taxa sampled in this study, including the corresponding GenBank accessions and sources for each species’ mitochondrial genome.

| **Order** | **Family** | **Species** | **Accession** | **Reference** |
| --- | --- | --- | --- | --- |
| Gonorhynchiformes | Gonorhynchidae | *Gonorhynchus abbeviatus* | AP009402 | (Lavoué et al., 2008) |
| Characiformes | Alestidae | *Phenacogrammus interruptus* | AB054129 | (Saitoh et al., 2003) |
| Characiformes | Bryconidae | *Brycon henni* | KP027535 | (Landínez-García et al., 2016) |
| Characiformes | Crenuchidae | *Crenuchus spilurus* | AP011986 | (Nakatani et al., 2011) |
| Characiformes | Serrasalmidae | *Serrasalmus eigenmanni* | PP754848 | (Condachou et al., 2024) |
| Characiformes | Acestrorhamphidae | *Astyanax mexicanus* | AP011982 | (Nakatani et al., 2011) |
| Characiformes | Lebiasinidae | *Nannostomus beckfordi* | OR857846 | (Xu et al., 2024) |
| Characiformes | Prochilodontidae | *Prochilodus argenteus* | KR014816 | (Chagas et al., 2016) |
| Cithariniformes | Citharinidae | *Citharinus congicus* | AP011985 | (Nakatani et al., 2011) |
| Cypriniformes | Botiidae | *Botia dario* | MN867911 | Unpublished |
| Cypriniformes | Catostomidae | *Catostomus commersonii* | AB127394 | (Saitoh et al., 2006) |
| Cypriniformes | Cyprinidae | *Cyprinus carpio* | OL699932 | Unpublished |
| Cypriniformes | Danionidae | *Danio rerio* | AC024175 | Unpublished |
| Cypriniformes | Leuciscidae | *Dionda episcopa* | AP012077 | (Iwasaki et al., 2013) |
| Gymnotiformes | Apteronotidae | *Apteronotus rostratus* | MH399592 | (Aguilar et al., 2019) |
| Gymnotiformes | Gymnotidae | *Electrophorus electricus* | AP011978 | (Nakatani et al., 2011) |
| Gymnotiformes | Hypopomidae | *Brachyhypopomus verdii* | KX058570 | (Elbassiouny et al., 2016) |
| Gymnotiformes | Rhamphichthyidae | *Gymnoramphichthys sp.* | AP011980 | (Nakatani et al., 2011) |
| Gymnotiformes | Sternopygidae | *Sternopygus dariensis* | MH399590 | (Aguilar et al., 2019) |
| Siluriformes | Ailiidae | *Ailia coila* | MK348534 | (Alam et al., 2019) |
| Siluriformes | Amblycipitidae | *Liobagrus styani* | KX096605 | (Huang et al., 2017) |
| Siluriformes | Amblycipitidae | *Proliobagrus dorsalis* | MN308285 | Unpublished |
| Siluriformes | Amphiliidae | *Amphilius* sp. | AP012002 | (Nakatani et al., 2011) |
| Siluriformes | Amphiliidae | *Zaireichthys* sp. | MZ930094 | (Schedel et al., 2022) |
| Siluriformes | Ariidae | *Ariopsis seemanni* | AP012003 | (Nakatani et al., 2011) |
| Siluriformes | Ariidae | *Arius arius* | KX211965 | Unpublished |
| Siluriformes | Ariidae | *Bagre panamensis* | KY930718 | (Ramírez-Pérez et al., 2017) |
| Siluriformes | Ariidae | *Bleekeriella leptaspis* | BK064997 | (Pimentel et al., 2023) |
| Siluriformes | Ariidae | *Brustiarius utarus* | BK065001 | (Pimentel et al., 2023) |
| Siluriformes | Ariidae | *Genidens barbus* | PP032974 | (Alvarenga et al., 2024) |
| Siluriformes | Ariidae | *Neoarius berneyi* | BK064999 | (Pimentel et al., 2023) |
| Siluriformes | Ariidae | *Occidentarius platypogon* | KY930717 | (Llera-Herrera et al., 2017) |
| Siluriformes | Ariidae | *Osteogeneiosus militaris* | MW566786 | Unpublished |
| Siluriformes | Aspredinidae | *Bunocephalus coracoideus* | AP012006 | (Nakatani et al., 2011) |
| Siluriformes | Astroblepidae | *Astroblepus* sp. | AP012004 | (Nakatani et al., 2011) |
| Siluriformes | Auchenipteridae | *Ageneiosus pardalis* | KM983421 | (Restrepo-Escobar et al., 2016) |
| Siluriformes | Auchenipteridae | *Duringlanis perugiae* | AP012024 | (Nakatani et al., 2011) |
| Siluriformes | Auchenipteridae | *Tetranematichthys quadrifilis* | AP012025 | (Nakatani et al., 2011) |
| Siluriformes | Auchenoglanididae | *Auchenoglanis occidentalis* | AP012005 | (Nakatani et al., 2011) |
| Siluriformes | Auchenoglanididae | *Parauchenoglanis monkei* | MZ930110 | (Schedel et al., 2022) |
| Siluriformes | Austroglanididae | *Austroglanis gilli* | MZ930072 | (Schedel et al., 2022) |
| Siluriformes | Austroglanididae | *Austroglanis sclateri* | MZ930070 | (Schedel et al., 2022) |
| Siluriformes | Bagridae | *Bagroides melapterus* | PV075036 | Unpublished |
| Siluriformes | Bagridae | *Bagrus caeruleus* | MZ930116 | (Schedel et al., 2022) |
| Siluriformes | Bagridae | *Bagrus docmak* | MZ930113 | (Schedel et al., 2022) |
| Siluriformes | Bagridae | *Hemibagrus guttatus* | PQ867598 | Unpublished |
| Siluriformes | Bagridae | *Mystus vittatus* | KX177968 | Unpublished |
| Siluriformes | Bagridae | *Sperata aor* | KX950699 | Unpublished |
| Siluriformes | Bagridae | *Tachysurus argentivittatus* | MW046162 | Unpublished |
| Siluriformes | Bagridae | *Hemibagrus wyckioides* | MF083116 | Unpublished |
| Siluriformes | Callichthyidae | *Brochis multiradiatus* | MN641874 | Unpublished |
| Siluriformes | Callichthyidae | *Dianema longibarbis* | PP737535 | Unpublished |
| Siluriformes | Callichthyidae | *Gastrodermus pygmaeus* | ON729306 | Unpublished |
| Siluriformes | Callichthyidae | *Hoplisoma concolor* | OQ569933 | Unpublished |
| Siluriformes | Callichthyidae | *Megalechis thoracata* | PP439705 | Unpublished |
| Siluriformes | Callichthyidae | *Osteogaster aenea* | MZ571336 | (Sun et al., 2022) |
| Siluriformes | Callichthyidae | *Scleromystax barbatus* | OQ507211 | (Dalcin et al., 2023) |
| Siluriformes | Cetopsidae | *Cetopsidium* sp. | AP012007 | (Nakatani et al., 2011) |
| Siluriformes | Cetopsidae | *Cetopsis coecutiens* | MW367090 | Unpublished |
| Siluriformes | Cetopsidae | *Helogenes marmoratus* | AP012014 | (Nakatani et al., 2011) |
| Siluriformes | Chacidae | *Chaca bankanensis* | AP012008 | (Nakatani et al., 2011) |
| Siluriformes | Clariidae | *Clarias batrachus* | NC_023923 | Unpublished |
| Siluriformes | Clariidae | *Clarias camerunensis* | OP936082 | Unpublished |
| Siluriformes | Clariidae | *Clarias dussumieri* | MG644387 | Unpublished |
| Siluriformes | Claroteidae | *Chrysichthys auratus* | MZ930111 | (Schedel et al., 2022) |
| Siluriformes | Claroteidae | *Chrysichthys* cf. *cranchii* | MZ930117 | (Schedel et al., 2022) |
| Siluriformes | Claroteidae | *Chrysichthys nigrodigitatus* | MH709123 | (Kim et al., 2018) |
| Siluriformes | Cranoglanididae | *Cranoglanis bouderius* | AY898626 | (Peng et al., 2006) |
| Siluriformes | Diplomystidae | *Diplomystes nahuelbutaensis* | AP012011 | (Nakatani et al., 2011) |
| Siluriformes | Doradidae | *Amblydoras gonzalezi* | AP012001 | (Nakatani et al., 2011) |
| Siluriformes | Doradidae | *Platydoras armatulus* | KM576101 | (Liu et al., 2016) |
| Siluriformes | Heptapteridae | *Brachyglanis microphthalmus* | PV662386 | (Arroyave et al., 2025) |
| Siluriformes | Heptapteridae | *Heptapterus mustelinus* | PV662388 | (Arroyave et al., 2025) |
| Siluriformes | Heptapteridae | *Imparfinis minutus* | PV662389 | (Arroyave et al., 2025) |
| Siluriformes | Heptapteridae | *Pimelodella montana* | PV662390 | (Arroyave et al., 2025) |
| Siluriformes | Heptapteridae | *Rhamdia guatemalensis* | PV662365 | (Arroyave et al., 2025) |
| Siluriformes | Heptapteridae | *Rhamdia laticauda* | PV662377 | (Arroyave et al., 2025) |
| Siluriformes | Heteropneustidae | *Heteropneustes fossilis* | AP012013 | (Nakatani et al., 2011) |
| Siluriformes | Horabagridae | *Horabagrus brachysoma* | KU870467 | Unpublished |
| Siluriformes | Horabagridae | *Horabagrus nigricollaris* | MG986722 | Unpublished |
| Siluriformes | Ictaluridae | *Ameiurus catus* | MG570433 | (Schroeter et al., 2020) |
| Siluriformes | Ictaluridae | *Ameiurus melas* | OM736826 | Unpublished |
| Siluriformes | Ictaluridae | *Ameiurus natalis* | MF621735 | (Schroeter et al., 2020) |
| Siluriformes | Ictaluridae | *Ameiurus nebulosus* | MF621733 | (Schroeter et al., 2020) |
| Siluriformes | Ictaluridae | *Ictalurus furcatus* | KM576102 | Unpublished |
| Siluriformes | Ictaluridae | *Ictalurus pricei* | KJ496298 | (Ballesteros-Córdova et al., 2016) |
| Siluriformes | Ictaluridae | *Ictalurus punctatus* | MF621716 | (Schroeter et al., 2020) |
| Siluriformes | Ictaluridae | *Noturus albater* | OR492264 | Unpublished |
| Siluriformes | Ictaluridae | *Noturus baileyi* | MW057778 | (Aunins et al., 2022) |
| Siluriformes | Ictaluridae | *Noturus exilis* | OR002151 | Unpublished |
| Siluriformes | Ictaluridae | *Noturus gyrinus* | OR492262 | Unpublished |
| Siluriformes | Ictaluridae | *Noturus insignis* | PQ064517 | Unpublished |
| Siluriformes | Ictaluridae | *Noturus leptacanthus* | OM736865 | Unpublished |
| Siluriformes | Ictaluridae | *Noturus taylori* | KP013089 | Unpublished |
| Siluriformes | Ictaluridae | *Prietella phreatophila* | MZ151888 | Unpublished |
| Siluriformes | Ictaluridae | *Pylodictis olivaris* | MF621730 | (Schroeter et al., 2020) |
| Siluriformes | Ictaluridae | *Trogloglanis pattersoni* | MZ151889 | Unpublished |
| **Siluriformes** | **Lacantuniidae** | ***Lacantunia enigmatica*** | **PX522361** | **This study** |
| Siluriformes | Loricariidae | *Ancistomus snethlageae* | KX087166 | (Moreira et al., 2017) |
| Siluriformes | Loricariidae | *Ancistrus cryptophthalmus* | MF804392 | (Mendes et al., 2019) |
| Siluriformes | Loricariidae | *Aphanotorulus emarginatus* | KT239019 | (Moreira et al., 2017) |
| Siluriformes | Loricariidae | *Hisonotus thayeri* | KX087173 | (Moreira et al., 2017) |
| Siluriformes | Loricariidae | *Hypoptopoma incognitum* | KT033767 | (Moreira et al., 2016) |
| Siluriformes | Loricariidae | *Hypostomus francisci* | MK026008 | (Pereira et al., 2019) |
| Siluriformes | Loricariidae | *Loricaria cataphracta* | KX087174 | (Moreira et al., 2017) |
| Siluriformes | Loricariidae | *Loricariichthys castaneus* | KT239015 | (Moreira et al., 2017) |
| Siluriformes | Loricariidae | *Pterygoplichthys pardalis* | KT239016 | (Moreira et al., 2017) |
| Siluriformes | Malapteruridae | *Malapterurus electricus* | OL802922 | Unpublished |
| Siluriformes | Mochokidae | *Atopochilus savorgnani* | MZ930077 | (Schedel et al., 2022) |
| Siluriformes | Mochokidae | *Atopodontus adriaensi* | MZ930079 | (Schedel et al., 2022) |
| Siluriformes | Mochokidae | *Chiloglanis* sp. 'Nigeria' | MZ930075 | (Schedel et al., 2022) |
| Siluriformes | Mochokidae | *Chiloglanis* sp. 'Bas Congo' | MZ930082 | (Schedel et al., 2022) |
| Siluriformes | Mochokidae | *Euchilichthys* sp. Kinsuka | MZ930102 | (Schedel et al., 2022) |
| Siluriformes | Mochokidae | *Microsynodontis* cf. *batesii* | MZ930109 | (Schedel et al., 2022) |
| Siluriformes | Mochokidae | *Microsynodontis* sp. 'Inga' | MZ930104 | (Schedel et al., 2022) |
| Siluriformes | Mochokidae | *Mochokiella paynei* | MZ930089 | (Schedel et al., 2022) |
| Siluriformes | Mochokidae | *Synodontis clarias* | OL450422 | Unpublished |
| Siluriformes | Mochokidae | *Synodontis petricola* | MZ930090 | (Schedel et al., 2022) |
| Siluriformes | Mochokidae | *Synodontis schoutedeni* | AP012023 | (Nakatani et al., 2011) |
| Siluriformes | Pangasidae | *Pangasianodon gigas* | AY762971 | Unpublished |
| Siluriformes | Pangasidae | *Pangasianodon hypophthalmus* | MZ272452 | (Duong et al., 2023) |
| Siluriformes | Pangasidae | *Pangasius bocourti* | MN842723 | Unpublished |
| Siluriformes | Pangasidae | *Pangasius conchophilus* | PP727124 | Unpublished |
| Siluriformes | Pangasidae | *Pangasius krempfi* | MZ272453 | (Duong et al., 2023) |
| Siluriformes | Pimelodidae | *Brachyplatystoma filamentosum* | PQ380008 | (Lilian Dantas Cavalcante et al., 2025) |
| Siluriformes | Pimelodidae | *Pimelodus pictus* | AP006768 | Unpublished |
| Siluriformes | Pimelodidae | *Pseudoplatystoma reticulatum* | KU291530 | (Villela et al., 2017) |
| Siluriformes | Pimelodidae | *Sorubim lima* | KY747394 | (Ren & Ma, 2019) |
| Siluriformes | Plotosidae | *Neosilurus hyrtlii* | OP882667 | Unpublished |
| Siluriformes | Plotosidae | *Plotosus japonicus* | AP012020 | (Nakatani et al., 2011) |
| Siluriformes | Plotosidae | *Porochilus rendahli* | OR263621 | Unpublished |
| Siluriformes | Plotosidae | *Tandanus tandanus* | OP882680 | Unpublished |
| Siluriformes | Pseudopimelodidae | *Lophiosilurus alexandri* | KJ494387 | (Carvalho et al., 2016) |
| Siluriformes | Pseudopimelodidae | *Pseudopimelodus schultzi* | MW959699 | (Yang & Ma, 2021) |
| Siluriformes | Ritidae | *Rita rita* | KF670723 | (Lashari et al., 2016) |
| Siluriformes | Schilbeidae | *Pareutropius debauwi* | AP012017 | (Nakatani et al., 2011) |
| Siluriformes | Schilbeidae | *Schilbe* cf. *intermedius* | MZ930118 | (Schedel et al., 2022) |
| Siluriformes | Schilbeidae | *Schilbe* cf. *yangambianus* | MZ930120 | (Schedel et al., 2022) |
| Siluriformes | Schilbeidae | *Schilbe grenfelli* | MZ930119 | (Schedel et al., 2022) |
| Siluriformes | Siluridae | *Kryptopterus bicirrhis* | KY569440 | Unpublished |
| Siluriformes | Siluridae | *Ompok bimaculatus* | KY887474 | (Barman et al., 2017) |
| Siluriformes | Siluridae | *Pterocryptis anomala* | MT433099 | (Chen et al., 2021) |
| Siluriformes | Siluridae | *Silurus asotus* | JX256247 | Unpublished |
| Siluriformes | Siluridae | *Wallago attu* | MN895039 | Unpublished |
| Siluriformes | Sisoridae | *Bagarius bagarius* | KJ204285 | (Lashari et al., 2016) |
| Siluriformes | Sisoridae | *Barbeuchiloglanis feae* | MZ901209 | (Schedel et al., 2022) |
| Siluriformes | Sisoridae | *Chimarrichthys davidi* | MK181572 | (Zou et al., 2019) |
| Siluriformes | Sisoridae | *Creteuchiloglanis macropterus* | KP872683 | (Ma et al., 2015) |
| Siluriformes | Sisoridae | *Erethistes jerdoni* | AP012012 | (Nakatani et al., 2011) |
| Siluriformes | Sisoridae | *Exostoma gaoligongense* | MW256713 | Unpublished |
| Siluriformes | Sisoridae | *Gagata dolichonema* | JQ026250 | Unpublished |
| Siluriformes | Sisoridae | *Glaridoglanis andersonii* | OK646330 | Unpublished |
| Siluriformes | Sisoridae | *Glyptosternon maculatum* | JQ026251 | Unpublished |
| Siluriformes | Sisoridae | *Glyptothorax annandalei* | MN396887 | Unpublished |
| Siluriformes | Sisoridae | *Pseudexostoma yunnanense* | JQ026258 | Unpublished |
| Siluriformes | Trichomycteridae | *Trichomycterus areolatus* | AP012026 | (Nakatani et al., 2011) |

**References**

Aguilar, C., Miller, M.J., Loaiza, J.R., Krahe, R., De León, L.F., 2019. Mitogenomics of Central American weakly-electric fishes. Gene. 686, 164–170. https://doi.org/10.1016/j.gene.2018.11.045.

Alam, M.J., Andriyono, S., Lee, S.R., Hossain, M.A.R., Eunus, A.T.M., Hassan, M.T., Kim, H.-W., 2019. Characterization of the complete mitochondrial genome of Gangetic ailia, *Ailia coila* (Siluriformes: Ailiidae). Mitochondrial DNA B Resour 4, 2258–2259. https://doi.org/10.1080/23802359.2019.1627942.

Alvarenga, M., D’Elia, A.K.P., Rocha, G., Arantes, C.A., Henning, F., de Vasconcelos, A.T.R., Solé-Cava, A.M., 2024. Mitochondrial genome structure and composition in 70 fishes: a key resource for fisheries management in the South Atlantic. BMC Genomics. 25, 215. https://doi.org/10.1186/s12864-024-10035-5.

Arroyave, J., Mar-Silva, A.F., Melo, B.F., Hernández-Ávila, S.G., López-Vila, J.M., Silva, G.S.C., Díaz-Jáimes, P., 2025. Evolutionary history of Middle American *Rhamdia* (Siluriformes: Heptapteridae) inferred from comparative mitogenomic data: Insights on historical biogeography and cave colonization in the group. J. Syst. Evol. 63, 1501-1518. https://doi.org/10.1111/jse.70003.

Aunins, A.W., Eackles, M.S., Super, P.E., Kulp, M.A., Nichols, B.J., Lubinski, B.A., Morrison, C.L., King, T.L., 2022. Development of a ddPCR assay for the detection of the Smoky Madtom (*Noturus baileyi*) from eDNA in stream water samples. Conservation Genet Resour. 14, 429–435. https://doi.org/10.1007/s12686-022-01290-3.

Ballesteros-Córdova, C.A., Castañeda-Rivera, M., Grijalva-Chon, J.M., Castillo-Gámez, R.A., Gutiérrez-Millán, L.E., Camarena-Rosales, F., Ruíz-Campos, G., Varela-Romero, A., 2016. Complete mitochondrial genome of *Ictalurus pricei* (Teleostei: Ictaluridae) and evidence of a cryptic *Ictalurus* species in Northwest Mexico. Mitochondrial DNA A DNA Mapp Seq Anal. 27, 4439–4441. https://doi.org/10.3109/19401736.2015.1089561.

Barman, A.S., Singh, M., Pandey, P.K., 2017. Complete mitochondrial genome of near threatened butter Catfish *Ompok bimaculatus* (Siluriformes: Siluridae). Mitochondrial DNA B Resour. 2, 313–314. https://doi.org/10.1080/23802359.2017.1334520.

Carvalho, D.C., Perini, V. da R., Bastos, A.S., Costa, I.R. da, Luz, R.K., Furtado, C., Prosdocimi, F., 2016. The complete mitochondrial genome of the threatened Neotropical catfish *Lophiosilurus alexandri* (Silurifomes: Pseudopimelodidae) and phylogenomic analysis indicate monophyly of Pimelodoidea. Genet Mol Biol. 39, 674–677. https://doi.org/10.1590/1678-4685-GMB-2016-0007.

Chagas, A.T. de A., Carmo, A.O., Costa, M.A., Resende, L.C., Brandão Dias, P.F.P., Martins, A.P.V., Kalapothakis, E., 2016. Description and comparison of two economically important fish species mitogenomes: *Prochilodus argenteus* and *Prochilodus costatus* (Characiformes, Prochilodontidae). Mitochondrial DNA A DNA Mapp Seq Anal. 27, 2852–2853. https://doi.org/10.3109/19401736.2015.1053125.

Chen, W., Li, Y., He, Y., Li, X., Li, J., 2021. Characterization of two complete mitochondrial genomes of *Pterocryptis anomala* (Siluridae) and its phylogeny and cryptic diversity. Biologia 76, 613–621. https://doi.org/10.2478/s11756-020-00582-z.

Condachou, C., Cuenot, Y., Pigeyre, L., Covain, R., Vigouroux, R., Brosse, S., Murienne, J., 2024. Genomic resources for the monitoring and management of *Tometes trilobatus*, *Hoplias aimara* and *Myloplus rhomboidalis*, three exploited freshwater fish species in French Guiana. Knowl. Manag. Aquat. Ecosyst. 17. https://doi.org/10.1051/kmae/2024011.

Dalcin, R.H., Ossa-Guerra, L.E.D. la, Artoni, R.F., Abilhoa, V., 2023. Complete mitochondrial genome of four *Scleromystax barbatus* (Siluriformes: Callichthyidae) populations. Neotrop. Ichthyol. 21, e230025. https://doi.org/10.1590/1982-0224-2023-0025.

Duong, T.Y., Pham, L.T.K., Le, X.T.K., Nguyen, N.T.T., Nor, A.M., Le, T.H., 2023. Mitophylogeny of Pangasiid Catfishes and its Taxonomic Implications for Pangasiidae and the Suborder Siluroidei. Zool Stud. 62, e48. https://doi.org/10.6620/ZS.2023.62-48.

Elbassiouny, A.A., Schott, R.K., Waddell, J.C., Kolmann, M.A., Lehmberg, E.S., Van Nynatten, A., Crampton, W.G.R., Chang, B.S.W., Lovejoy, N.R., 2016. Mitochondrial genomes of the South American electric knifefishes (Order Gymnotiformes). Mitochondrial DNA B Resour. 1, 401–403. https://doi.org/10.1080/23802359.2016.1174090.

Huang, J.-Y., Hu, S., Bai, X., Zhang, E., 2017. Complete mitochondrial genome of *Liobagrus styani* (Teleostei: Amblycipitidae). Mitochondrial DNA B Resour. 2, 15–16. https://doi.org/10.1080/23802359.2016.1275841.

Iwasaki, W., Fukunaga, T., Isagozawa, R., Yamada, K., Maeda, Y., Satoh, T.P., Sado, T., Mabuchi, K., Takeshima, H., Miya, M., Nishida, M., 2013. MitoFish and MitoAnnotator: A Mitochondrial Genome Database of Fish with an Accurate and Automatic Annotation Pipeline. Mol. Biol. Evol. 30, 2531–2540. https://doi.org/10.1093/molbev/mst141.

Kim, N.-K., Zealous Gietbong, F., Andriyono, S., Kim, A.R., Kim, H.-W., 2018. The complete mitogenome of Bagrid catfish *Chrysichthys nigrodigitatus* (Siluriformes: Claroteidae). Mitochondrial DNA B Resour. 3, 1239–1240. https://doi.org/10.1080/23802359.2018.1532341.

Landínez-García, R.M., Alzate, J.F., Márquez, E.J., 2016. Complete mitogenome of the Neotropical fish *Brycon henni*, Eigenmann 1913 (Characiformes, Bryconidae). Mitochondrial DNA A DNA Mapp Seq Anal 27, 2259–2260. https://doi.org/10.3109/19401736.2014.984170.

Lashari, P., Laghari, M.Y., Xu, P., Zhao, Z., Jiang, L., Narejo, N.T., Deng, Y., Sun, X., Zhang, Y., 2016. Complete mitochondrial genome of catfish *Bagarius bagarius* (Hamilton, Sisoridae; Siluriformes) from Indus River Sindh, Pakistan. Mitochondrial DNA A DNA Mapp Seq Anal. 27, 439–440. https://doi.org/10.3109/19401736.2014.900612.

Lavoué, S., Miya, M., Poulsen, J.Y., Møller, P.R., Nishida, M., 2008. Monophyly, phylogenetic position and inter-familial relationships of the Alepocephaliformes (Teleostei) based on whole mitogenome sequences. Mol. Phylogenet. Evol. 47, 1111–1121. https://doi.org/10.1016/j.ympev.2007.12.002.

Lilian Dantas Cavalcante, R., Santos Silva, C., Ferreira Vidal, A., Soares Pires, É., Lopes Nunes, G., Fogaça de Assis Montag, L., Oliveira, G., Ribeiro-dos-Santos, Â., Santos, S., José de Souza, S., Estefano de Santana Souza, J., Sakamoto, T., 2025. The complete mitogenome of Amazonian *Brachyplatystoma filamentosum* and the evolutionary history of body size in the order Siluriformes. Sci. Rep. 15, 9873. https://doi.org/10.1038/s41598-025-94272-y.

Liu, S., Yao, J., Zhang, J., Liu, Z., 2016. Next generation sequencing yields the complete mitochondrial genome of the striped raphael catfish, *Platydoras armatulus* (Siluriformes: Doradidae). Mitochondrial DNA A DNA Mapp. Seq. Anal. 27, 1963–1964. https://doi.org/10.3109/19401736.2014.971308.

Llera-Herrera, R., Ramírez-Pérez, J.S., Saavedra-Sotelo, N.C., 2017. Complete mitochondrial genome of Cominate sea catfish *Occidentarius platypogon* (Siluriformes: Ariidae). Mitochondrial DNA B Resour. 2, 337–338. https://doi.org/10.1080/23802359.2017.1334516.

Ma, X., Kang, J., Chen, W., Zhou, C., He, S., 2015. Biogeographic history and high-elevation adaptations inferred from the mitochondrial genome of Glyptosternoid fishes (Sisoridae, Siluriformes) from the southeastern Tibetan Plateau. BMC Evol. Biol. 15, 233. https://doi.org/10.1186/s12862-015-0516-9.

Mendes, I.S., Prosdocimi, F., Schomaker-Bastos, A., Furtado, C., Ferreira, R.L., Santos Pompeu, P., Carvalho, D.C., 2019. On the evolutionary origin of Neotropical cavefish *Ancistrus cryptophthalmus* (Siluriformes, Loricariidae) based on the mitogenome and genetic structure of cave and surface populations. Hydrobiologia 842, 157–171. https://doi.org/10.1007/s10750-019-04033-y.

Moreira, D.A., Buckup, P.A., Furtado, C., Val, A.L., Schama, R., Parente, T.E., 2017. Reducing the information gap on Loricarioidei (Siluriformes) mitochondrial genomics. BMC Genomics 18, 345. https://doi.org/10.1186/s12864-017-3709-3.

Moreira, D.A., Magalhães, M.G.P., de Andrade, P.C.C., Furtado, C., Val, A.L., Parente, T.E., 2016. An RNA-based approach to sequence the mitogenome of *Hypoptopoma incognitum* (Siluriformes: Loricariidae). Mitochondrial DNA A DNA Mapp. Seq. Anal. 27, 3784–3786. https://doi.org/10.3109/19401736.2015.1079903.

Nakatani, M., Miya, M., Mabuchi, K., Saitoh, K., Nishida, M., 2011. Evolutionary history of Otophysi (Teleostei), a major clade of the modern freshwater fishes: Pangaean origin and Mesozoic radiation. BMC Evol. Biol. 11, 177. https://doi.org/10.1186/1471-2148-11-177.

Peng, Z., Wang, J., He, S., 2006. The complete mitochondrial genome of the helmet catfish *Cranoglanis bouderius* (Siluriformes: Cranoglanididae) and the phylogeny of otophysan fishes. Gene. 376, 290–297. https://doi.org/10.1016/j.gene.2006.04.014.

Pereira, A.H., Facchin, S., Oliveira do Carmo, A., Núñez Rodriguez, D., Cardoso Resende, L., Kalapothakis, Y., Brandão Dias Ferreira Pinto, P., Mascarenhas Alves, C.B., Henrique Zawadzki, C., Kalapothakis, E., 2019. Complete mitochondrial genome sequence of *Hypostomus francisci* (Siluriformes: Loricariidae). Mitochondrial DNA B Resour. 4, 155–157. https://doi.org/10.1080/23802359.2018.1544860.

Pimentel, L.G.P., Silva, I.B. da, Rodrigues-Oliveira, I.H., Pasa, R., Menegídio, F.B., Kavalco, K.F., 2023. Description of eight new mitochondrial genomes for the genus *Neoarius* and phylogenetic considerations for the family Ariidae (*Siluriformes*). Genomics Inform. 21. https://doi.org/10.5808/gi.23059.

Ramírez-Pérez, J.S., Saavedra-Sotelo, N.C., Llera-Herrera, R., Abadía-Chanona, Q.Y., 2017. Complete mitochondrial genome of the Chihuil sea catfish *Bagre panamensis* (Siluriformes: Ariidae). Mitochondrial Mitochondrial DNA B Resour. 2, 341–343. https://doi.org/10.1080/23802359.2017.1334519.

Ren, F., Ma, X., 2019. The complete mitochondrial genome of *Sorubim Lima* (Siluriformes, Pimelodidae). Mitochondrial DNA B Resour. 4, 3650–3651. https://doi.org/10.1080/23802359.2019.1678420.

Restrepo-Escobar, N., Alzate, J.F., Márquez, E.J., 2016. Mitochondrial genome of the Neotropical catfish *Ageneiosus pardalis*, Lütken 1874 (Siluriformes, Auchenipteridae). Mitochondrial DNA A DNA Mapp. Seq. Anal. 27, 2176–2177. https://doi.org/10.3109/19401736.2014.982613.

Saitoh, K., Miya, M., Inoue, J.G., Ishiguro, N.B., Nishida, M., 2003. Mitochondrial Genomics of Ostariophysan Fishes: Perspectives on Phylogeny and Biogeography. J. Mol. Evol. 56, 464–472. https://doi.org/10.1007/s00239-002-2417-y.

Saitoh, K., Sado, T., Mayden, R.L., Hanzawa, N., Nakamura, K., Nishida, M., Miya, M., 2006. Mitogenomic evolution and interrelationships of the Cypriniformes (Actinopterygii: Ostariophysi): the first evidence toward resolution of higher-level relationships of the world’s largest freshwater fish clade based on 59 whole mitogenome sequences. J. Mol. Evol. 63, 826–841. https://doi.org/10.1007/s00239-005-0293-y.

Schedel, F.D.B., Chakona, A., Sidlauskas, B.L., Popoola, M.O., Usimesa Wingi, N., Neumann, D., Vreven, E.J.W.M.N., Schliewen, U.K., 2022. New phylogenetic insights into the African catfish families Mochokidae and Austroglanididae. J. Fish Biol. 100, 1171–1186. https://doi.org/10.1111/jfb.15014.

Schroeter, J.C., Maloy, A.P., Rees, C.B., Bartron, M.L., 2020. Fish mitochondrial genome sequencing: expanding genetic resources to support species detection and biodiversity monitoring using environmental DNA. Conserv. Genet. Resour. 12, 433–446. https://doi.org/10.1007/s12686-019-01111-0.

Sun, C.-H., Huang, Q., Zeng, X.-S., Li, S., Zhang, X.-L., Zhang, Y.-N., Liao, J., Lu, C.-H., Han, B.-P., Zhang, Q., 2022. Comparative analysis of the mitogenomes of two *Corydoras* (Siluriformes, Loricarioidei) with nine known *Corydoras*, and a phylogenetic analysis of Loricarioidei. ZooKeys. 1083, 89–107. https://doi.org/10.3897/zookeys.1083.76887.

Villela, L.C.V., Alves, A.L., Varela, E.S., Yamagishi, M.E.B., Giachetto, P.F., da Silva, N.M.A., Ponzetto, J.M., Paiva, S.R., Caetano, A.R., 2017. Complete mitochondrial genome from South American catfish *Pseudoplatystoma reticulatum* (Eigenmann & Eigenmann) and its impact in Siluriformes phylogenetic tree. Genetica 145, 51–66. https://doi.org/10.1007/s10709-016-9945-7.

Xu, W., Tai, J., He, K., Xu, T., Zhang, G., Xu, B., Liu, H., 2024. Complete Mitochondrial Genomes of *Nannostomus* Pencilfish: Genome Characterization and Phylogenetic Analysis. Animals 14, 1598. https://doi.org/10.3390/ani14111598.

Yang, H., Ma, X., 2021. The complete mitochondrial genome of *Pseudopimelodus schultzi* Dahl 1955 (Siluriformes, Pseudopimelodidae) and its phylogenetic position within Pseudopimelodidae. Mitochondrial Mitochondrial DNA B Resour. 6, 2206–2208. https://doi.org/10.1080/23802359.2021.1945974.

Zou, Y., Hu, H., Zhang, P., Wen, Z., Wei, Q., 2019. The complete mitochondrial genome of *Euchiloglanis davidi* and its phylogenetic implications. Mitochondrial Mitochondrial DNA B Resour. 4, 1249–1250. https://doi.org/10.1080/23802359.2019.1566789.
