## Supplementary Table 2 for "African origin and Late Cretaceous divergence of the Middle American catfish *Lacantunia enigmatica* corroborated by a global mitogenome phylogeny"

**Supplementary Table 2.** Genes and associated features of the mitochondrial genome of *Lacantunia enigmatica*. Intergenic space (IGS) described as intergenic (+) or overlapping nucleotides (–).

| **Locus** | **Type** | **One-letter code** | **Start** | **End** | **Length (bp)** | **Strand** | **# of AA** | **Anticodon** | **Start codon** | **Stop codon** | **IGS** |
| --- | --- | --- | --- | --- | --- | --- | --- | --- | --- | --- | --- |
| *tRNA^Phe^* | tRNA | F | 1 | 69 | 69 | H |  | GAA |  |  | 0 |
| *12s rRNA* | rRNA |  | 70 | 1017 | 948 | H |  |  |  |  | 0 |
| *tRNA-Val* | tRNA | V | 1018 | 1091 | 74 | H |  | TAC |  |  | 0 |
| *16s rRNA* | rRNA |  | 1092 | 2766 | 1092 | H |  |  |  |  | 0 |
| *tRNA-Leu* | tRNA | L | 2767 | 2840 | 74 | H |  | TAA |  |  | 63 |
| *NAD1* | Protein-coding |  | 2904 | 3872 | 951 | H | 316 |  | ATG | TAA | 7 |
| *tRNA^Ile^* | tRNA | I | 3880 | 3949 | 70 | H |  | GAT |  |  | 8 |
| *tRNA^Gln^* | tRNA | Q | 3958 | 4028 | 71 | L |  | TTG |  |  | –1 |
| *tRNA^Met^* | tRNA | M | 4028 | 4097 | 70 | H |  | CAT |  |  | 0 |
| *NAD2* | Protein-coding |  | 4098 | 5144 | 1047 | H | 337 |  | ATG | AGA | –3 |
| *tRNA^Trp^* | tRNA | W | 5142 | 5211 | 70 | H |  | TCA |  |  | 1 |
| *tRNA^Ala^* | tRNA | A | 5213 | 5281 | 69 | L |  | TGC |  |  | 1 |
| *tRNA^Asn^* | tRNA | N | 5283 | 5355 | 73 | L |  | GTT |  |  | 53 |
| *tRNA^Cys^* | tRNA | C | 5409 | 5475 | 67 | L |  | GCA |  |  | 0 |
| *tRNA^Tyr^* | tRNA | Y | 5476 | 5542 | 67 | L |  | GTA |  |  | 1 |
| *COX1* | Protein-coding |  | 5544 | 7131 | 1528 | H | 489 |  | GTG | AGA | –4 |
| *tRNA^Ser^* | tRNA | S | 7127 | 7197 | 71 | L |  | TGA |  |  | 2 |
| *tRNA^Asp^* | tRNA | D | 7200 | 7270 | 71 | H |  | GTC |  |  | 2 |
| *COX2* | Protein-coding |  | 7273 | 7963 | 691 | H | 225 |  | ATG | T | 0 |
| *tRNA^Lys^* | tRNA | K | 7964 | 8036 | 73 | H |  | TTT |  |  | 1 |
| *ATP8* | Protein-coding |  | 8038 | 8205 | 168 | H | 51 |  | ATG | TAA | –8 |
| *ATP6* | Protein-coding |  | 8196 | 8878 | 683 | H | 223 |  | ATG | TA | 0 |
| *COX3* | Protein-coding |  | 8879 | 9662 | 784 | H | 249 |  | ATG | T | 0 |
| *tRNA^Gly^* | tRNA | G | 9663 | 9731 | 69 | H |  | TCC |  |  | 0 |
| *NAD3* | Protein-coding |  | 9732 | 10079 | 348 | H | 112 |  | ATG | GAC | 0 |
| *tRNA^Arg^* | tRNA | R | 10080 | 10148 | 69 | H |  | TCG |  |  | 0 |
| *NAD4L* | Protein-coding |  | 10149 | 10445 | 297 | H | 97 |  | ATA | TAA | –5 |
| *NAD4* | Protein-coding |  | 10439 | 11819 | 1380 | H | 445 |  | ATG | T | 0 |
| *tRNA^His^* | tRNA | H | 11820 | 11888 | 69 | H |  | GTG |  |  | 0 |
| *tRNA^Ser^* | tRNA | S | 11889 | 11952 | 64 | H |  | GCT |  |  | –1 |
| *tRNA^Leu^* | tRNA | L | 11952 | 12024 | 73 | H |  | TAG |  |  | 1 |
| *NAD5* | Protein-coding |  | 12026 | 13855 | 1830 | H | 598 |  | ATG | TA | –2 |
| *NAD6* | Protein-coding |  | 13852 | 14373 | 522 | L | 172 |  | ATG | T | 1 |
| *tRNA^Glu^* | tRNA | E | 14375 | 14443 | 69 | L |  | TTC |  |  | 2 |
| *CYB* | Protein-coding |  | 14446 | 15586 | 1141 | H | 369 |  | ATG | T | 0 |
| *tRNA^Thr^* | tRNA | T | 15587 | 15662 | 76 | H |  | TGT |  |  | –1 |
| *tRNA-Pro* | tRNA | P | 15662 | 15730 | 69 | L |  | TGG |  |  | 0 |
| *D-loop* | Non-coding |  | 15731 | 16804 | 1074 | H |  |  |  |  | 0 |
